# Plasticity after cortical stroke involves potentiating responses of pre-existing circuits but not functional remapping to new circuits

**DOI:** 10.1101/2020.11.09.375840

**Authors:** William A Zeiger, Máté Marosi, Satvir Saggi, Natalie Noble, Isa Samad, Carlos Portera-Cailliau

## Abstract

Functional recovery after stroke is thought to be mediated by adaptive circuit plasticity, whereby surviving neurons assume the roles of those that died. This “remapping” hypothesis is based on human brain mapping studies showing apparent reorganization of cortical sensorimotor maps and animal studies documenting molecular and structural changes that could support circuit rewiring. However, definitive evidence of remapping is lacking, and other studies have suggested that maladaptive plasticity mechanisms, such as enhanced inhibition in peri-infarct cortex, might actually limit plasticity after stroke. Here we sought to directly test whether neurons can change their response selectivity after a stroke that destroys a single barrel (C1) within mouse primary somatosensory cortex. Using multimodal in vivo imaging approaches, including two-photon calcium imaging to longitudinally record sensory-evoked activity in peri-infarct cortex before and after stroke, we found no evidence to support the remapping hypothesis. In an attempt to promote plasticity via rehabilitation, we also tested the effects of forced use therapy by plucking all whiskers except the C1 whisker. Again, we failed to detect an increase in the number of C1 whisker-responsive neurons in surrounding barrels even 2 months after stroke. Instead, we found that forced use therapy potentiated sensory-evoked responses in a pool of surviving neurons that were already C1 whisker responsive by significantly increasing the reliability of their responses. Together, our results argue against the long-held theory of functional remapping after stroke, but support a plausible circuit-based mechanism for how rehabilitation may improve recovery of function.

## Introduction

Stroke is the fifth leading cause of death and the leading cause of adult-onset disability in the U.S. (Benjamin et al., 2018). Many stroke patients exhibit partial spontaneous recovery of function, which can be improved with rehabilitation (Corbetta et al., 2015; Pollock et al., 2014). This suggests that the brain has endogenous mechanisms to restore lost functions, a process known as neuroplasticity. The prevailing hypothesis is that post-stroke plasticity involves a process of circuit remapping in which neurons that survive the injury are somehow recruited to assume new responsibilities to take over the role of neurons that died (Carmichael, 2012; Murphy and Corbett, 2009; Xerri et al., 2014). Certainly, functional remapping occurs in the healthy developing brain during learning (Barth and Ray, 2019) and in response to changes in sensory experience (Hooks and Chen, 2020; Margolis et al., 2014). However, in the context of stroke, the available data on circuit plasticity has not yet provided irrefutable evidence for remapping of lost functionalities to new circuits.

For example, human studies with brain imaging or neurophysiology have revealed variable (and sometimes opposite) changes in brain activity or metabolism after stroke. Some of these changes, such as the apparent reorganization of cortical sensorimotor maps (Altamura et al., 2007; Carey et al., 2011; Cramer et al., 1997; Delvaux et al., 2003; Jang, 2011; Jang et al., 2005; Rossini et al., 1998; Schaechter et al., 2006; Zemke et al., 2003), or differences in resting state functional connectivity (Bannister et al., 2015; Dijkhuizen et al., 2014; Goodin et al., 2018) have been interpreted as evidence of remapping. But they could also reflect normal variability across individuals, especially when considering that these imaging modalities could be influenced by altered hemodynamics post-stroke and that pre-stroke baseline data was not available.

In mice, macroscopic brain imaging studies have similarly shown a range of alterations in cortical activity maps after stroke (Brown et al., 2009; Harrison et al., 2013; Jablonka et al., 2010; Kraft et al., 2018; Mohajerani et al., 2011), but whether those were associated with functional recovery has not been established. In a landmark in vivo calcium imaging study often cited as the best evidence for remapping after stroke, some neurons in peri-infarct cortex were more broadly tuned after stroke compared to neurons in control mice without a stroke (Winship et al., 2008). However, the fraction of neurons that had assumed new roles was rather small and, just like with the human studies, circuits were not imaged longitudinally before and after stroke.

Another intriguing issue regarding stroke plasticity is that several studies have identified circuit alterations that would be expected to limit rather than promote remapping. In human stroke, compensation by the unaffected limb appears to induce maladaptive plasticity (Jones, 2017; Takeuchi and Izumi, 2012), perhaps via increased interhemispheric inhibition (Boddington and Reynolds, 2017). Pathological increases in inhibition after stroke have also been identified in animal models of stroke, and reducing this inhibition can promote recovery (Alia et al., 2016; Clarkson et al., 2010). Other studies have found that experience-dependent plasticity in both somatosensory and visual cortex is impaired following strokes targeted to the somatosensory cortex (Greifzu et al., 2011; Jablonka et al., 2007, 2012). In summary, definitive evidence for functional remapping after stroke remains lacking, and whether all aspects of post-stroke plasticity are beneficial remains controversial.

Addressing these knowledge gaps about stroke plasticity requires longitudinal in vivo recordings of the activity of individual neurons before and after stroke. Toward that end, we performed in vivo intrinsic signal imaging (ISI) and two-photon (2P) calcium imaging of sensory-evoked responses before and after a photothrombotic (PT) stroke that was targeted to a single barrel in the barrel field of primary somatosensory cortex (S1BF). We found no clear evidence of remapping of lost functionalities to new circuits in peri-infarct cortex. However, plucking all whiskers except the one corresponding to the infarcted barrel (as a forced use rehabilitative strategy) significantly enhanced the reliability of responses in neuronal ensembles that were already responsive to that whisker at baseline. Our results argue against the classic remapping model of stroke recovery where surviving neurons/circuits can assume a new role. Rather, our data is more consistent with a model where spontaneous or rehabilitation-induced recovery involves potentiation of pre-existing circuits.

## Materials & Methods

### Materials

All chemicals were obtained from Sigma Aldrich unless otherwise noted.

#### Experimental Animals

All experiments followed the U.S. National Institutes of Health guidelines for animal research, under an animal use protocol approved by the Chancellor’s Animal Research Committee (ARC) and Office for Animal Research Oversight at the University of California, Los Angeles (#2005-145). Both male and female adult mice were used. All animals were housed in a vivarium with a 12 h light/dark cycle. For in vivo imaging, transgenic *Thy1-GCaMP6s* mice (GP4.3, JAX line 024275) (Dana et al., 2014) were used. For activity-dependent labeling, we used the TRAP (<u>T</u>argeted <u>R</u>ecombination in <u>A</u>ctive <u>P</u>opulations) approach (Guenthner et al., 2013), crossing *cFos*-CreER^T2^ mice (JAX line 021882) with the Ai9 Cre-dependent tdTomato reporter line (JAX line 007909). All transgenic lines were maintained on a C57BL/J6 background.

#### Cranial Window Surgery

We implanted chronic glass-covered cranial windows in 6-10 week old mice as described previously (Holtmaat et al., 2009; Mostany and Portera-Cailliau, 2008). Animals were deeply anesthetized using 5% isoflurane followed by maintenance with 1.5-2% isoflurane. A circular craniotomy, ~4-5 mm in diameter, was made using a pneumatic dental drill over the primary somatosensory cortex, centered ~3 mm lateral to the midline and ~2 mm caudal to Bregma. A sterile glass coverslip was placed over the craniotomy and glued to the skull with cyanoacrylate glue (Krazy Glue). Dental acrylic (OrthoJet, Lang Dental) was then applied throughout the exposed skull surface around the edges of the coverslip. To later secure the mouse onto the microscope stage, a small titanium bar was embedded in the dental acrylic. Carprofen (5 mg/kg, i.p., Zoetis) and dexamethasone (0.2 mg/kg, i.p., Vet One) were provided for pain relief and mitigation of edema on the day of surgery and daily for the next 48 h. Mice were allowed to recover from the surgery for 2-4 weeks before the first imaging session.

#### Intrinsic Signal Imaging

Whisker-evoked sensory activity maps were generated using intrinsic signal imaging (ISI) as previously described (Johnston et al., 2013). Animals were sedated with chlorprothixene (~3 mg/kg, i.p.), head-fixed to the stage of a custom-built tandem-lens macroscope, and lightly anesthetized with ~0.5-0.7% isoflurane. The cortical surface was illuminated by green (535 nm) LEDs to visualize the superficial vasculature. The macroscope was then focused ~300 μm below the cortical surface and red (630 nm) LEDs were used to record intrinsic signals, with frames collected 0.9 s before and 1.5 s after stimulation by a fast CCD camera at 30 Hz (Teledyne Dalse Pantera 1M60). Thirty trials separated by 20 s were conducted for each imaging session. Whisker stimuli (100 Hz, 1.5 s duration) were delivered by carefully affixing an individual whisker to a glass capillary tube coupled to a ceramic piezoelectric bending actuator (Physik Instrumente). Evoked signals were quantified as the mean pixel intensity of the evoked whisker map normalized to the background signal outside of the map. If a map could not be visualized (for instance, following stroke), the mean pixel intensity of the area adjacent to the stroke was normalized to the background signal. For qualitative assessment of whisker-evoked ISI maps, raw ISI signals were manually thresholded to create a binary mask. The mask was then applied to the raw ISI signal and the signal over threshold was overlaid on the cortical vasculature image. To determine the number of animals in which a C1-evoked ISI map was present at 2 months post-stroke, ISI images were each scored for presence or absence of a map based on consensus of three experimenters blinded to group assignment.

#### Photothrombotic Stroke

Animals were deeply anesthetized as above, and core body temperature was maintained at 37°C using a homeothermic blanket system (Harvard Apparatus). Rose Bengal dye (120 mg/kg of mouse body weight, diluted in sterile saline, i.p.), was injected 10 min prior to head-fixing the mouse on the stage of a custom-built two photon microscope. The beam of a green laser (532 nm, ~2 mW intensity at the sample; Laserlands 1875-532D) was aligned through the optical path of the microscope and scanned across an ~0.5 x 0.5 mm region of the cortical surface corresponding to the C1-barrel (previously identified using ISI) for 10 min. Animals were allowed to recover on a heated water-recirculating blanket in the dark for 1 h before being returned to the home cage. For the sham control group, animals were injected with an equivalent volume of saline, but otherwise treated the same, with laser scanning across the C1 barrel.

#### Cresyl Violet Staining

Five days after stroke animals were transcardially perfused with 4% paraformaldehyde in phosphate buffered saline. Brains were removed and 50 μm thick coronal sections were cut using a vibrating microtome (Leica VT 1000). Sections were sequentially mounted, dried, and placed into 1:1 ethanol:chloroform overnight. The following day, sections were rehydrated through 100% and 95% ethanol, rinsed in dH2O, and then stained in 0.1% cresyl violet solution (warmed to 37°C) for 10 minutes. After staining, sections were rinsed in dH2O, differentiated in 95% ethanol for 10 minutes, dehydrated in 100% ethanol, and cleared in xylene. Sections were imaged on an upright fluorescent microscope (Zeiss Axio Imager 2) using a 10x objective and transmitted light.

#### Whisker Trimming/Plucking

After animals were appropriately anesthetized (as described above), all whiskers on the right side of the snout, except those undergoing stimulation (C1 and/or D3), were trimmed using a fine scissors to a length of ~5 mm on the morning prior to imaging. For the forced use whisker plucking group, all whiskers on the right side of the face except C1 were entirely plucked from the follicle starting 24 h after stroke. Additional plucking was performed three times weekly to remove any whisker re-growth. After one month, whiskers were allowed to re-grow. For sham plucking, mice were similarly anesthetized, and we handled their whiskers with forceps but did not actually remove the whisker.

#### Two-photon In Vivo Calcium Imaging

Calcium imaging was performed as previously described using a custom-built two-photon microscope with galvanometric scanners (Cambridge Technology), a Chameleon Ultra II Ti:sapphire laser (Coherent), a 20x objective (0.95 NA, Olympus), and ScanImage software (Vidrio Technologies) (He et al., 2017). Data for 3/18 mice in the forced use experiment (one control animal at one month, one plucked animal at one month, and one control animal at two months post-stroke) were collected on a galvo-resonant microscope (Neurolabware) due to a hardware failure on the primary microscope. Mice were lightly sedated with chlorprothixene (~3 mg/kg, i.p.) and isoflurane (0.7-0.9%) and kept warm with a heating blanket. Stimulation of the C1 or D3 whiskers (20 stimuli, 1 s duration at 10 Hz, with a 3 s interstimulus interval) was delivered with a piezoelectric actuator, as above. Whole-field images were acquired with bidirectional scanning at ~7.8 Hz (1024 x 128 pixels, down-sampled to 256 x 128 pixels). On average, we recorded 31.8 active neurons per field-of-view (FOV). In total, we recorded from 13,991 neurons in 33 mice. For each imaging field-of-view (FOV), ~100 s of spontaneous activity data was collected prior to sensory-evoked activity.

Fluorescence traces (*ΔF/F*) of neuronal calcium transients were extracted using custom-written semi-automated MATLAB routines as previously described (He et al., 2017). The vast majority of movies (>90%) did not require motion correction, but for those that did, *X–Y* drift was corrected using either a frame-by-frame, hidden Markov model–based registration routine (Dombeck et al., 2007) or the motion correction module from *EZcalcium* (Cantu et al., 2020) based on *NoRMCorre* nonrigid template matching (Pnevmatikakis and Giovannucci, 2017). To determine whether individual neurons showed time-locked responses to whisker stimulations, we first calculated a modified *Z* score vector for each neuron and then used a probabilistic bootstrapping method to correlate calcium transients with epochs of stimulation, as described (He et al., 2017).

For spontaneous activity, amplitude and frequency of calcium transients were detected using the PeakFinder script in MATLAB. Sensory-evoked activity from cells was quantified in several ways. Cells were separated into groups based on whether they exhibited time-locked responses to whisker stimulation (time-locked or non-time-locked). Modified Z-scores for each of the 20 individual whisker stimuli were aligned to the onset of the stimulus and a mean stimulus-evoked trace was calculated for each cell. The area-under-the-curve (AUC) of the mean stimulus-evoked trace was quantified as the trapezoidal integral of the modified Z-score vector over 2 s after stimulus onset. In this way, the AUC of the mean stimulus-evoked trace captures the stimulus-evoked activity of a cell across all 20 stimuli. For each neuron we also used the PeakFinder script to identify the presence (or absence) of an evoked response, the peak amplitude of that response, and the latency to peak for each of the 20 individual whisker stimuli delivered in a given imaging session. For peak amplitude and latency, the mean response was calculated only for stimulus epochs with a detected peak; stimulus epochs without a detected peak were ignored. Because many neurons show decreases in the amplitude of their calcium signals with ongoing bouts of whisker stimulation (He et al., 2017), we also calculated adaptation indices across the 20 stimuli as:

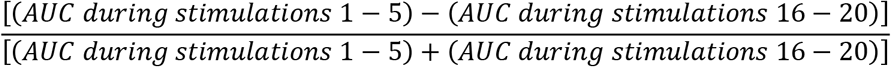

#### Activity-dependent labeling

Transgenic mouse lines, in which the Cre recombinase is knocked in downstream of the promoter for the immediate early gene *cFos,* have recently been developed to enable activity-dependent labeling of neurons during a defined epoch (Guenthner et al., 2013). This approach has been termed Targeted Recombination in Active Populations, or TRAP. We crossed these *cFos*-CreER^T2^ mice with the Cre-dependent reporter line Ai9. TRAP mice underwent cranial window implantation and stroke (or sham stroke) of the C1 barrel. ISI imaging was used first to locate the C1 barrel and then again one week after stroke to confirm proper targeting. Two months after stroke, mice were brought to the behavior testing room and all whiskers were trimmed flush to the face, except for the right-sided C1 whisker. The following day, mice were injected with 4-hydroxytamoxifen (50 mg/kg in corn oil, i.p.) as previously described (Guenthner et al., 2013). Following injection, mice were placed in pairs in an environmental enrichment cage in the dark for 6 h. Mice were then returned to their home cages for 3-7 d, followed by transcardial perfusion with 4% paraformaldehyde in phosphate buffered saline.

Brains were removed and the left hemisphere was carefully dissected and cut into 60 μm thick coronal sections using a vibrating microtome (Leica VT 1000). Sections were sequentially mounted using Fluoromount-G with DAPI (ThermoFisher). The section containing the C1 barrel (for sham animals) or stroke was manually identified and 4 sections anterior and 2 sections posterior, totaling a span of ~360 μm in the anterior/posterior direction were imaged on an upright fluorescent microscope (Zeiss Axio Imager 2) using a 10x objective, Apotome.2 optical sectioning module, and DAPI or DsRed filter sets. Images were then randomized and blinded for quantification of tdTomato-labeled putative C1-whisker responsive cells in the peri-infarct cortex. The S1BF was first identified on each DAPI-labeled image and manually annotated using ImageJ and the mouse Allen Brain atlas as a reference. The region-of-interest corresponding to the S1BF was then transferred to the tdTomato images and all tdTomato-positive cells were manually counted, by cortical layer, within the S1BF region. Labeled cells within the ischemic core or the C1-barrel were excluded from analysis of peri-infarct labeled cells.

#### Statistical Analyses

All data are shown as mean +/- standard error of the mean, unless otherwise stated. Sample sizes were not based on *a priori* power calculations but are consistent with other studies in the field using similar techniques, including our own (Goel et al., 2018; He et al., 2017; Johnston et al., 2013; Mostany et al., 2010). Statistical analyses were performed using MATLAB. Data were tested for normality using Lilliefors test and analyzed using parametric or non-parametric statistical tests as indicated in the figure legends.

For percentage of whisker-responsive cells (**Fig. 2C-D, 4C**), generalized linear mixed effects models were used with binomial distribution, according to the model “ ResponsiveROIs ~ 1 + Group + Timepoint + Timepoint*Group + (1 | MouseID)”, with each animal serving as one data point. In the analysis of percentage of whisker-responsive cells comparing control to whisker-plucked groups (**Fig. 4C**), we found no effects of group or group-by-time interaction, so data were pooled and re-analyzed just for effects of time according to the model “ ResponsiveROIs ~ 1 + Timepoint + (1 | MouseID)”.

For analysis of AUC, peak amplitude, and peak latency of sensory-evoked calcium events (**Fig 5E-F, s6**), linear mixed effects models on data from individual neurons were used, according to the formula “y ~ Group + Timepoint + Group * Timepoint + (1 | MouseID)”. Residuals from raw data were non-normal, so data was log-transformed for further statistical analysis. To facilitate log transformation, a constant value was added to any datasets with negative values, such that the minimum value for the dataset was equal to 0.1. For analysis of percentage of stimuli for which a neuron was active, a generalized linear mixed effects model was used with binomial distribution. For all linear mixed effects models, *p* values for individual coefficients from the model were corrected for multiple hypothesis testing using the Benjamini & Hochberg procedure (Benjamini and Hochberg, 1995). ANOVA was used to test for overall effects of group, time, and group by time interactions. In the figures, results are presented with *p* values for overall effects of group (G), timepoint (T) and group-by-timepoint interactions (G:T) shown as an inset on respective graphs, with *p* values for individual coefficients depicted adjacent to the respective data points.

This is the first longitudinal two-photon calcium imaging study to record neuronal activity for weeks before and after stroke, a technically challenging undertaking. In some animals, the cranial window did not remain optically transparent for every single imaging time point. As a result, data for some time points could not be collected. For the analysis of the percentage of neurons with time-locked response to C1 and D3 whisker stimulation, comparing sham versus stroke groups, the following data points were missing: one mouse in the stroke group at 13 d post-stroke and 4 mice in each group at one month post-stroke. For the analysis of sensory-evoked responses in control versus forced use whisker plucked groups, one mouse in the plucked group had no neurons with time-locked responses to C1 whisker stimulation at +13 d post-stroke.

For correlation between stroke size and percentage of time-locked cells or ISI signal, data was fit using a simple linear regression model. Values for r^2^ and *p* are presented on the graphs for pooled data. Analysis was also conducted on each group separately and no significant correlation was seen for either group individually.

## Results

To study neuronal plasticity and remapping at the local circuit level before and after focal cortical lesions, we focused on the barrel field of primary somatosensory cortex (S1BF). We chose S1BF for several reasons: 1. Somatosensory deficits are common after stroke in humans (Connell et al., 2008; Kessner Simon S. et al., 2019); 2. As nocturnal animals, mice preferentially rely on their whiskers to explore their surroundings; 3. S1BF exhibits significant experience-dependent plasticity, even in adult animals (Gainey and Feldman, 2017; Glazewski and Fox, 1996; Margolis et al., 2014); and 4. The dynamics of somatosensory circuit function in rodents are well characterized (Chen et al., 2015; Diamond and Arabzadeh, 2013; Peron et al., 2015; Petersen, 2019). The S1BF exhibits precise somatotopic organization with each barrel receiving inputs primarily from its corresponding peripheral whisker (labeled by row and arc position, e.g., C1, D3, etc.) (Diamond and Arabzadeh, 2013), but also containing neurons that respond to adjacent barrels (Clancy 2015).

To create focal cortical lesions, we adapted the PT stroke model to target a small region in S1BF, just slightly larger than an individual barrel, and largely sparing neighboring barrels, i.e., the peri-infarct region (**Fig. 1**). The resulting single barrel infarcts (**Fig. 1B**) were meant to mimic most strokes in humans, which tend to be very small, with an average infarct size of <5% of total brain volume, and frequently <1% (Brott et al., 1989; Fiebach et al., 2015; Hakimelahi et al., 2014; Lansberg et al., 2001; Sperber and Karnath, 2016). Importantly, our goal was to leave adjacent barrels largely intact, expecting that they would engage in compensatory plasticity. Larger strokes affecting the entire S1BF would presumably require involvement of more distant regions that are less homologous in function. Indeed, based on popular theories about plasticity (Murphy and Corbett, 2009), we hypothesized that spared barrels adjacent to a stroke in a single barrel would take over the lost functionality because they serve an analogous function for a different whisker. In addition, within a given barrel, individual neuronal responses are heterogenous, with a small but substantial population of neurons that are preferentially tuned not to the principal whisker of that barrel but to adjacent whiskers (Clancy et al., 2015). Furthermore, this whisker selectivity is highly dynamic and can adapt to changes in sensory experience (Margolis et al., 2012).

**Figure 1.**
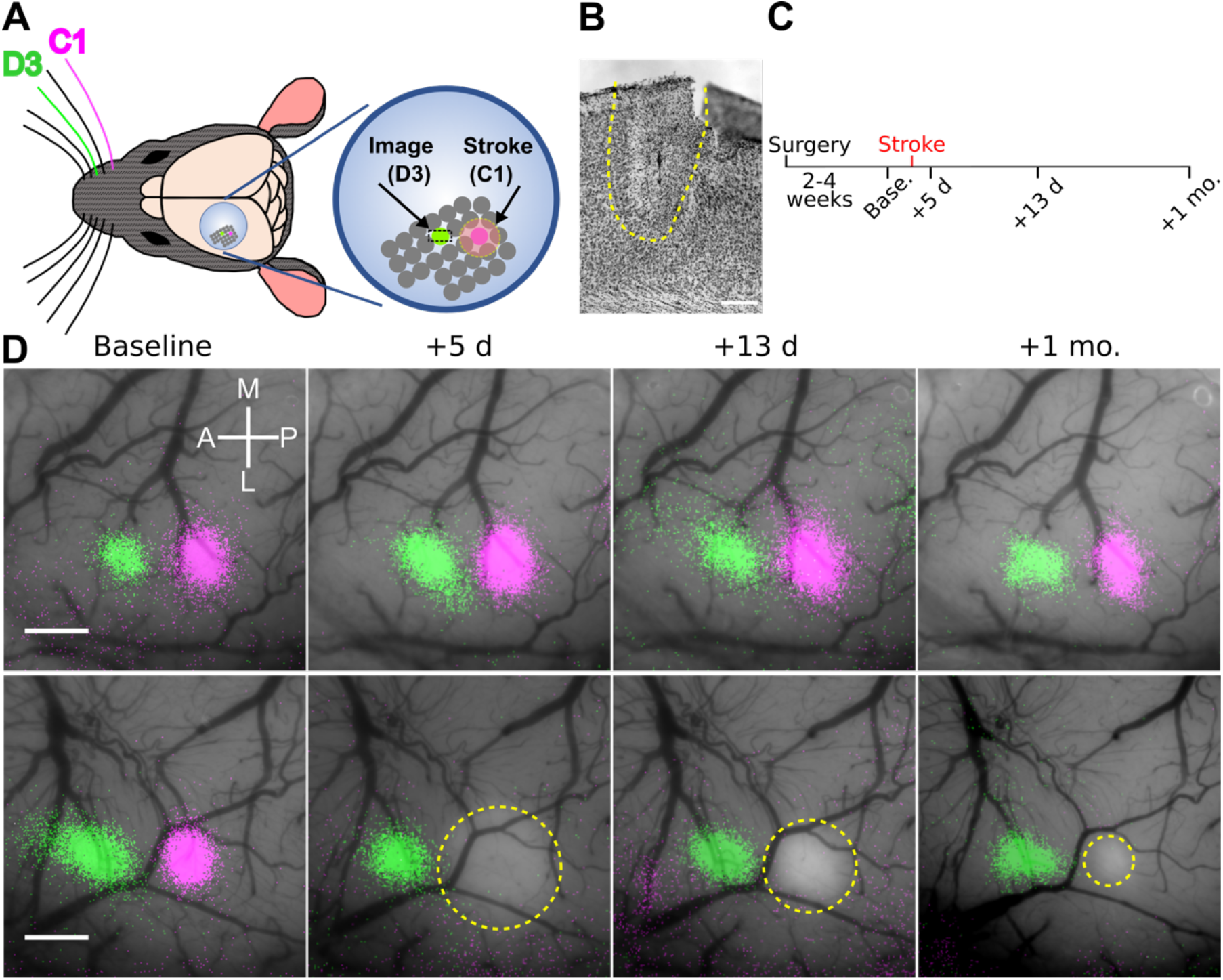
Intrinsic signal imaging reveals no evidence of macroscopic remapping after single barrel photothrombotic strokes. **A.** Left: Schematic of cranial window placement and whiskers for stimulation. Right: Enlarged view of cranial window with the locations of the infarcted barrel (C1, magenta) and spared barrel (D3, green) highlighted. **B.** Cresyl Violet stained coronal section from a representative mouse 5 days post-stroke; the infarct core is outlined in yellow. Scale bar = 200 μm. **C.** Timeline for surgery and imaging. ISI and 2P imaging were carried out at each of the indicated timepoints – baseline (before stroke) and +5 days, +13 days, and +1 month after stroke. **D.** ISI maps elicited by stimulation of the C1 (magenta) and D3 (green) whiskers overlaid on photographs of the cranial window in representative sham control (top) and stroke (bottom) mice before and after stroke. Pale areas (yellow outline) represent the infarct. Note that the infarct size appears smaller over time as a result of the disappearance of acute tissue edema and tissue involution and scarring. Scale bar = 0.5 mm.

For an initial set of experiments, we implanted cranial windows over S1BF in young adult mice and targeted PT strokes to the C1 barrel (**Fig. 1**).We used ISI (Gao et al., 2017) to identify single whisker-evoked maps in S1BF at the pre-stroke baseline and longitudinally for over one month following stroke (**Fig, 1C**). Before stroke, we found that ISI activity maps for the C1 and D3 barrels were spatially distinct, with the D3 map located anterior to the C1 map, as expected (**Fig. 1D**). In sham control animals (exposed to laser light but not rose Bengal; see Methods), the C1 and D3 whisker-evoked maps remained stable in size and location throughout the duration of imaging (**Fig. 1D,** upper panel). In stroke animals, we could discern the PT infarct within the C1 barrel as a pale area where the ISI map was previously located (**Fig. 1D,** lower panel). Following stroke, D3 whisker-evoked activity maps remained unaffected, whereas the C1 whisker-evoked activity map disappeared and did not recover even one month after stroke (**Fig 1D,** lower panel).

There are several reasons why the C1 whisker-evoked ISI activity map might not re-emerge after stroke. One possibility is that plasticity simply does not occur after stroke because spared brain regions in peri-infarct cortex cannot compensate for lost functionalities. Another reason is related to technical limitations of ISI, which probably lacks the resolution to detect the activity of small numbers of neurons (scattered over a large cortical area or multiple layers) that exhibit compensatory plasticity (i.e., their signals are not sufficient to generate an ISI map). Alternatively, changes in blood flow and oxygen context after stroke (i.e., impaired neurovascular coupling (He et al., 2020)) may affect our ability to detect new ISI maps after stroke. To address these concerns about ISI, we also recorded whisker-evoked responses in the spared D3 barrel with in vivo 2P calcium imaging, which provides single cell resolution and is, in principle, impervious to changes in blood flow (**Fig. 2A-B)**.

**Figure 2.**
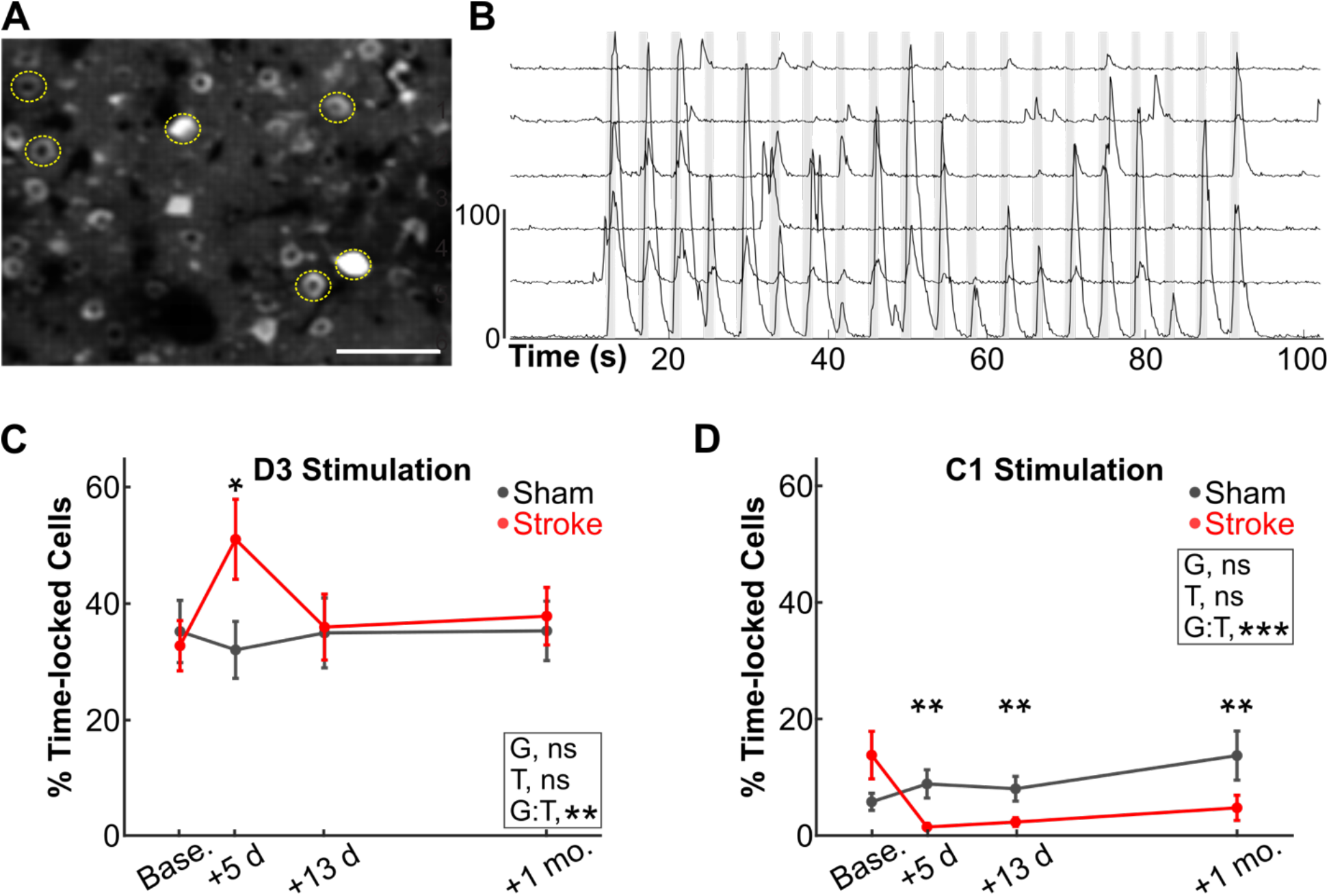
No increase in the percentage of C1 whisker-responsive neurons in peri-infarct cortex after C1 barrel stroke. **A.** Maximum intensity projection image of GCaMP6s fluorescence in the D3 barrel of a Thy1-GCaMP6s mouse (line 4.3), before stroke. Scale bar 100 μm. **B.** Representative traces of GCaMP6s fluorescence signal intensity from 6 active L2/3 neurons (corresponding to the somata circled in *A)* in response to whisker stimulation (10 Hz). Scale bar indicates Z-score change of 100. Vertical gray bars indicate epochs of whisker stimulation (1 s long, 3 s inter-stimulus interval). **C. D.** Percentage of L2/3 neurons in the D3 barrel showing responses that were time-locked to either D3 (C) or C1 (D) whisker stimulation for mice that received a sham procedure (gray; laser light but without rose Bengal, see Methods) or a photothrombotic stroke to the C1 barrel (red). Individual time points are means ± s.e.m. (n= 6 and 9 mice for sham and stroke, respectively). GLME binomial model, main effects of group (G), timepoint (T) and group-by-timepoint interaction (G:T). Significance for individual coefficients for G:T indicated over corresponding data points (*, *p*<0.05; **, *p*<0.01).

We recorded calcium signals of L2/3 neurons in S1BF of Thy1-GCaMP6s mice in response to 20 sequential stimulations of the C1 or D3 whiskers (see Methods; **Fig. 2B**). Before stroke, we found that, on average, 32.6 ± 4.3% of L2/3 neurons in the D3 barrel responded to stimulation of their principal D3 whisker with time-locked responses (**Fig. 2C**), consistent with our previous studies (He et al., 2017). In contrast, only 13.9 ± 4.1% of L2/3 neurons in the D3 barrel responded to the surrounding C1 whisker (**Fig. 2D**), consistent with published studies using calcium imaging (Clancy et al., 2015). As evidence of remapping of lost representation of the C1 whisker after stroke, we predicted that the percentage of C1-responsive neurons within the D3 barrel would increase after stroke. However, we found instead that the percentage of L2/3 neurons with calcium transients time-locked to stimulation of the C1 whisker actually decreased significantly to <2 % at 5 d post-stroke and remained below pre-stroke levels even up to one month post-stroke (**Fig. 2D**). Interestingly, the percentage of D3 barrel neurons responding to the D3 whisker was transiently (at 5 d post-stroke) higher than at baseline (**Fig. 2C**). In sham control animals, the percentage of C1 and D3 whisker-responsive cells remained stable across imaging sessions (**Fig. 2C-D**, black lines).

As a complementary approach to test the hypothesis that recovery and circuit remapping after a stroke that destroys the C1 barrel is mediated by the recruitment of new neurons that respond to the C1 whisker in surrounding barrels, we employed an activity-dependent labeling strategy using TRAP (Targeted Recombination in Active Populations). This approach was developed to label neurons activated by specific stimuli during defined epochs and has previously been used to label neurons responsive to stimulation of an individual whisker in vivo (Guenthner et al., 2013). We randomized double transgenic (cFos-CreER^T2^ x Ai9) TRAP mice to sham or PT strokes of the C1 barrel and allowed the animals to recover spontaneously for 2 months. Mice then underwent trimming of all whiskers except the contralesional C1 whisker. The next day, mice were injected with the CreER^T2^ ligand 4-OHT and allowed to explore an enriched environment in the dark for 6 h (see Methods). TRAP resulted in robust labeling of neurons in the C1 barrel in sham animals, primarily in L2/3 and L4 (**Fig. 3A**). Sparser labeling was also observed in other layers and in surrounding barrels, as expected. We next quantified the number of fluorescently labeled neurons in the surrounding peri-infarct S1BF, representing putative C1-whisker responsive neurons. Compared to sham control animals, we did not find an increase in the total number of C1-whisker responsive neurons in stroke mice (**Fig. 3B**), even when looking at individual cortical layers (**Suppl. Fig. 1**). Together, results from this initial set of experiments, which failed to detect a new C1 whisker map (by ISI) or an increase in the proportion of C1 whisker-responsive neurons (calcium imaging and TRAP), do not support the theory of remapping of neuronal function after focal cortical injury.

**Figure 3.**
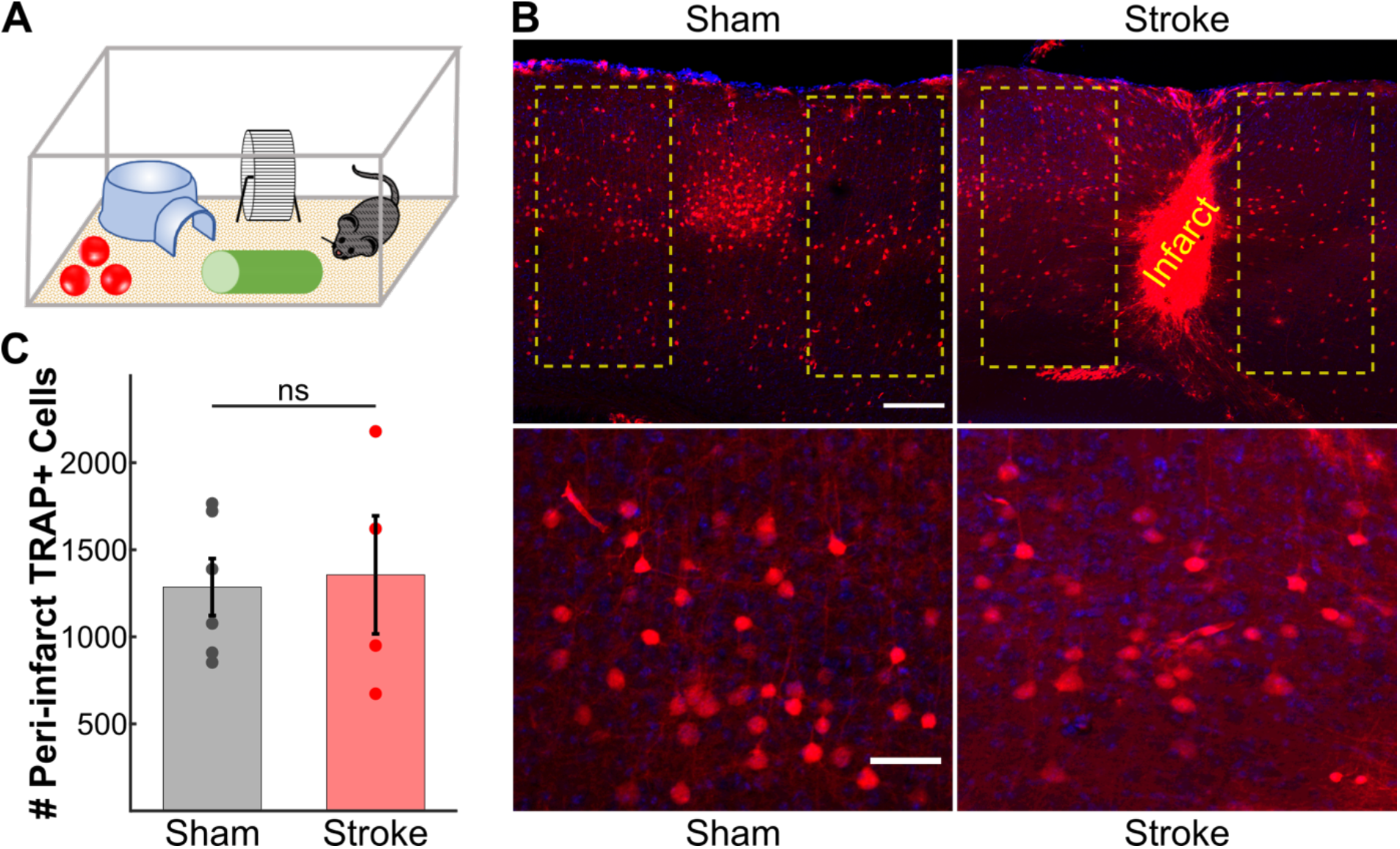
TRAP labeling shows no increase in the numbers of C1 whisker-responsive neurons in the peri-infarct cortex 2 months after stroke. **A.** Schematic of TRAP labeling approach. Two months after stroke targeting the C1 barrel, *cFos*-CreER^T2^:Ai9 mice were subjected to whisker trimming (all whisker except C1) and then allowed to explore an enriched environment using only the right sided C1 whisker for 6 hours immediately following injection of 4-OHT (50 mg/kg in corn oil). **B.** Top: Representative images of activity dependent TRAP labeling in S1BF. Putative C1-whisker responsive neurons are labeled in red (Td-Tom); DAPI counterstain. Example regions for quantification are outlined in yellow surrounding either the intact C1 barrel in sham control mice (left panel) or the infarct core in stroke mice (right panel). Note the large density of cFos-expressing (TRAP+) neurons in the C1 barrel in sham control mice. Scale bar = 200 μm. Bottom: High magnification of L4 TRAP labeled cells surrounding the intact C1 barrel in sham control mice (left panel) or the infarct core in stroke mice (right panel). Scale bar = 50 μm. **C.** Quantification of the total number of TRAP+ neurons in the peri-infarct cortex (stroke group) or surround barrels (sham group). Two-sample *t*-test, *p*=0.804. N= 6 and 4 sham and stroke mice, respectively.

It is possible that circuit remapping can only occur under certain circumstances in which additional plasticity mechanisms are engaged, like in the setting of rehabilitation (Cassidy and Cramer, 2017; Stinear et al., 2020). In a second set of experiments, we explored this possibility by utilizing whisker plucking, a well-established paradigm for inducing neuronal plasticity in the S1BF. Whisker plucking has been shown to promote circuit remapping in the intact S1BF through changes in dendritic spines (Holtmaat et al., 2006; Miquelajauregui et al., 2015; Schubert et al., 2013) or in the excitation/inhibition ratio (Gainey and Feldman, 2017; Li et al., 2014), via re-tuning of L2/3 neurons away from deprived whiskers and toward spared whiskers (Margolis et al., 2012), and by an expansion of whisker-evoked ISI activity maps (Gao et al., 2017; Margolis et al., 2012; Polley et al., 1999). We also favored this approach as it is analogous to a rehabilitation technique in humans known as forced use therapy. This therapy, whereby the “good” or unaffected limb is restrained in order to force the patient to use their affected limb, has been shown to promote recovery of function after stroke in humans (Corbetta et al., 2015; Kwakkel et al., 2015). Thus, we reasoned that forced use of the C1 whisker after stroke might boost inherent plasticity mechanisms and lead to an increase in the number of C1 whisker-responsive neurons in peri-infarct cortex, just as what happens after whisker deprivation in the intact S1BF.

Beginning ~24 h after stroke we plucked all whiskers on the contralesional side of the snout, except the C1 whisker (corresponding to the infarcted barrel) (**Fig. 4A-B**); we continued whisker plucking three times per week until one month after stroke. Control animals were subjected to a sham plucking procedure (see Methods). We again began by performing ISI to monitor recovery of C1 whisker-evoked activity maps and extended our imaging timepoints to two months post-stroke (**Fig. 4B-D**). The majority of mice in the control group (5/8) did not exhibit any recovery of the C1 map at 2 months post-stroke, although we did see smaller maps in 3/8 mice as early as 13 d post-stroke (**Fig 4C-D**). Qualitatively, a slightly greater proportion of animals in the whisker-plucked group (5/10) had a detectable C1 whisker-evoked ISI activity map at 2 months compared to the control group (**Fig. 4E**). However, quantification of C1 whisker-evoked intrinsic signal over the entire cranial window showed a decline in signal intensity at 5 d after stroke that remained lower than pre-stroke levels even at 2 months post-stroke, and there was no difference between control and whisker-plucked groups (**Fig. 4F**). Stroke size at 5 d post-stroke was similar between groups (**Fig. 4G)** and there was a trend toward an inverse correlation between stroke size and C1 whisker-evoked ISI signal at 2 months post-stroke (**Suppl. Fig. 2A**), suggesting the partial map recovery seen in some animals may be attributable to incomplete ablation of the entire C1 barrel.

**Figure 4.**
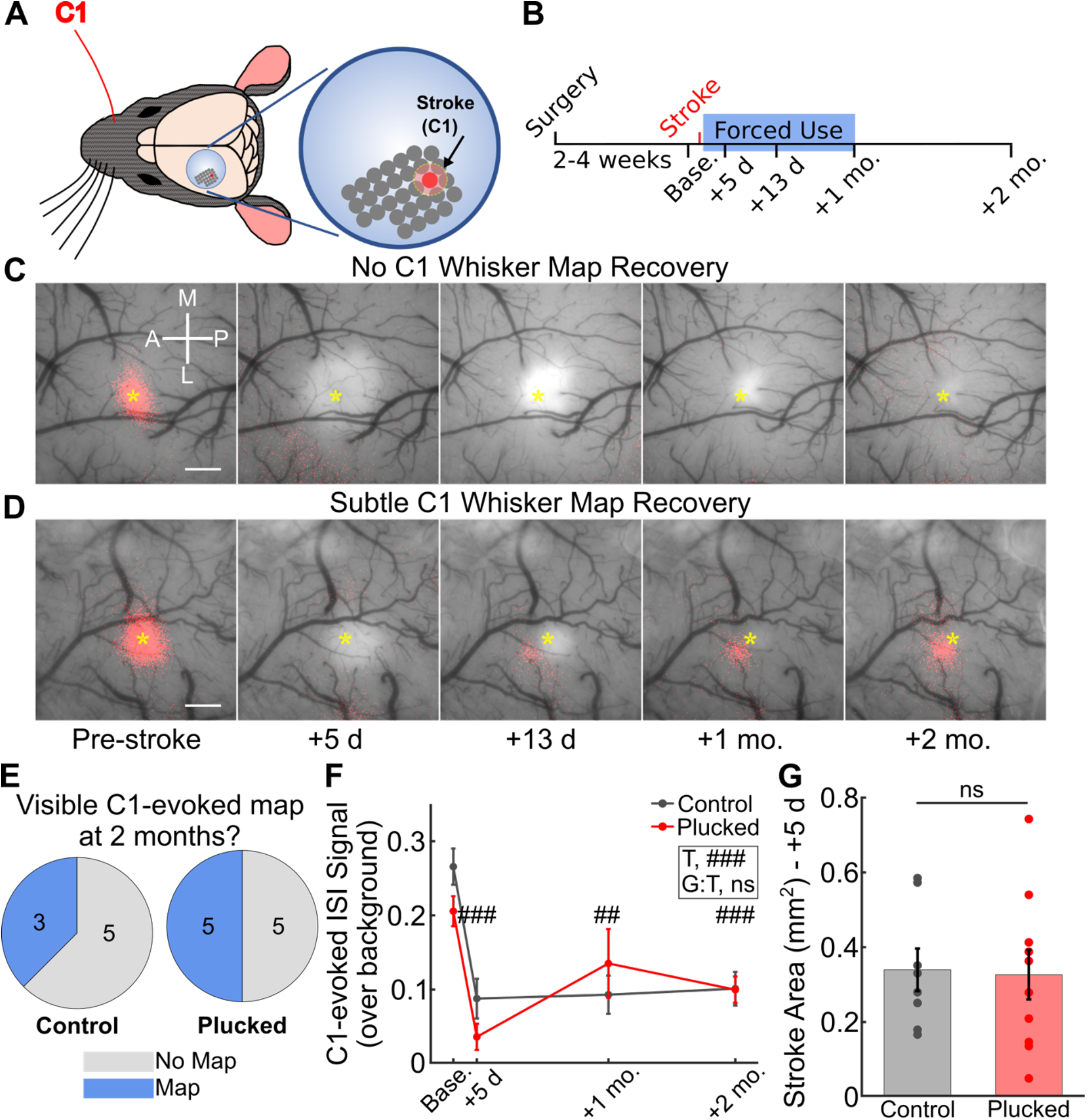
Forced use therapy does not trigger macroscopic remapping after single barrel stroke. **A.** Schematic of forced use rehabilitation paradigm, where all right-sided whiskers are plucked except C1. **B.** Timeline of surgery, imaging, and forced use whisker plucking. **C-D.** ISI map of C1 whisker-evoked activity before and after stroke overlaid on photographs of the cranial window in representative mice (with strokes in C1 barrel) that exhibited no recovery of the C1 whisker map (C) or subtle partial map recovery (D) over 2 months. Scale bar = 0.5 mm. **E.** Qualitative assessment of C1 whisker-evoked activity maps at 2 months after stroke in the control stroke group vs. the forced use group all with whisker plucked except C1 (χ^2^, *p*=0.596; n= 8 and 10, respectively). **F.** Quantification of C1 whisker-evoked ISI signals over background throughout recovery in control vs. forced-use (whisker plucked) animals (n= 8 and 10, respectively). Twoway repeated measures ANOVA, main effects of timepoint (T, ###, *p*<0.001) and group-by-timepoint interaction (G:T, *p*=0.182). Multiple comparison testing performed using the Tukey-Kramer procedure. Significance for individual coefficients for timepoints (T) indicated over corresponding data points (##, *p*<0.01; ###, *p*<.001). **G.** Quantification of stroke size at day 5 after stroke from vasculature images (see Methods). Individual points indicate different animals (n= 8 and 10 mice for control and plucked, respectively). Two-sample *t*-test, *p*=.888.

Next, we performed 2P calcium imaging of neuronal responses to C1 whisker stimulation in whisker plucked mice (forced use therapy). Here, we chose to image L2/3 neurons not just in the D3 barrel, but in multiple fields of view (FOV) across the peri-infarct region of S1BF (typically 2-3 FOV, with 32 active neurons on average per FOV) at each time point before and after stroke (**Fig. 5A**). First, we confirmed that, at baseline (before the stroke), the percentage of neurons with time-locked responses to C1 whisker stimulation within the C1 barrel was similar between whisker-plucked animals and the non-plucked control group (34.3 ± 2.6% and 34.8 ± 5.3%, respectively; **Fig. 5B**). Similarly, the percentage of neurons with time-locked responses to C1 whisker stimulation in FOVs adjacent to (but not overlapping) the C1 barrel was similar between plucked and control groups (14.1 ± 1.8% vs. 17.2 ± 3.1%, respectively; **Fig. 5C)**. Following stroke, we observed a slight decrease in the percentage of C1 time-locked neurons in both control and whisker-plucked groups at 5 d and 13 d post-stroke (**Fig. 5C**), similar to what we had observed in the D3 barrel in our initial cohort of animals (see **Fig. 2D**, red line). By 1 month and 2 months post-stroke, the percentage of neurons in the peri-infarct FOVs with time-locked responses to C1 whisker stimulation had recovered to pre-stroke levels in both groups, without significant differences between the groups (**Fig. 5C**). A similar trend was observed when normalizing each animal to its pre-stroke baseline percentage of neurons with time-locked responses (**Suppl. Fig. 3A**). In other words, the percentage of neurons in peri-infarct cortex that responded to the C1 whisker does not increase above baseline after stroke, even with rehabilitation, which means that the forced use therapy does not promote remapping in peri-infarct cortex. If anything, there is a transient decrease in the proportion of C1-whisker responsive neurons in peri-infarct cortex, consistent with previous reports of increased inhibition acutely post-stroke (Alia et al., 2016; Clarkson et al., 2010).

**Figure 5.**
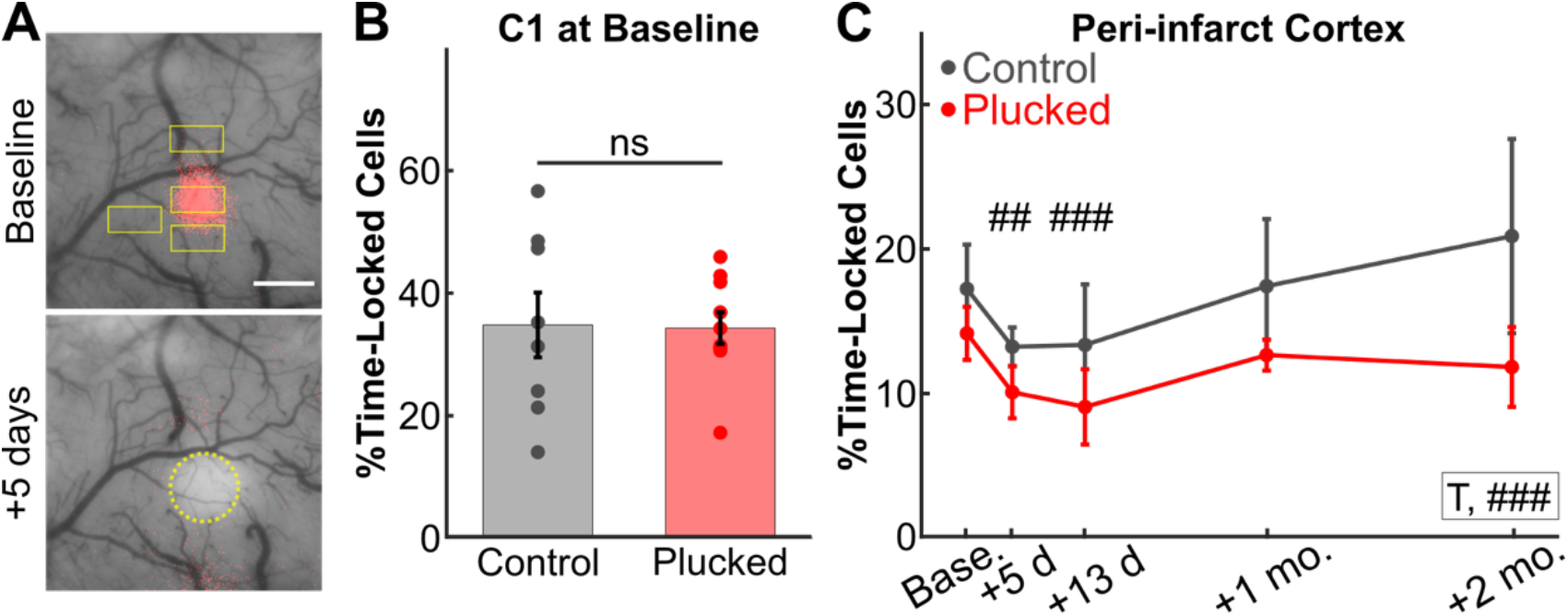
Forced use therapy does not lead to an increase in the number of whisker responsive cells in peri-infarct cortex. **A.** Representative ISI maps depicting C1 whisker-evoked activity (red) overlaid on a photograph of the cranial window at baseline (top) or 5 days post-stroke (bottom). Yellow rectangles indicate the locations of calcium imaging fields-of-view throughout the peri-infarct region. Stroke at +5 days outlined in yellow circle. **B.** Percentage of L2/3 neurons in the C1 barrel with time-locked responses to C1-whisker stimulation at baseline prior to stroke in control (all whiskers intact) or forced use groups (all whiskers plucked except C1) (Two-sample *t*-test, *p=* 0.929; N= 8 and 10 mice, respectively). **C.** Percentage of neurons with time-locked responses to C1-whisker stimulation in the peri-infarct regions throughout recovery in control and forced use groups. GLME binomial model with separate groups, main effects of group (G, *p*=0.338), timepoint (T, *p*=0.208), and group-by-timepoint interaction (G:T, *p=*0.554) were not significant. GLME binomial model with pooled groups, main effect of timepoint (T, ###, *p*=<0.001), with significance for individual coefficients for timepoints (T) indicated over corresponding data points (##, *p*<0.01; ###, *p*<0.001).

We then reasoned that, if circuit remapping was restricted to a highly localized area, we might potentially be diluting an effect by including neurons from multiple, potentially uninvolved FOVs. Therefore, we next focused our analysis on FOVs with the greatest percentage of C1 time-locked neurons at each timepoint. In this analysis, the FOV selected might differ at each timepoint, depending on which FOV has the highest proportion of C1 time-locked neurons. The trend was very similar to the results analyzing all FOVs, with a transient decrease at 5 d and 13 d post-stroke, followed by recovery to pre-stroke baseline levels, and no difference between the plucked and the control cohorts (**Suppl. Fig. 3B**). Even when plotting individual mice, we did not see any obvious differences between the two cohorts. We also longitudinally analyzed changes in the FOVs with the lowest percentage of C1 time-locked neurons at baseline (analyzing the same FOV for each mouse at each timepoint here), supposing that there might be a ceiling to the percentage of whisker time-locked neurons within a given barrel, which limits recruitment of those barrels for post-stroke plasticity. This analysis showed considerable variability but, once again, there was no increase in the percentage of C1 time-locked neurons after stroke and no differences between cohorts (**Suppl. Fig 3C**).

Overall, these results argue against the possibility that new neurons from peri-infarct cortex are recruited to subserve the functions of those lost after stroke. We also considered that the degree of potential plasticity at the level of individual neurons (i.e., the ability of neurons to change their tuning to the C1 whisker after stroke) is driven by the size of the stroke, such that, for instance, larger strokes might limit plasticity. However, we found that, although animals with larger infarcts at 5 d post-stroke tended to have fewer C1 time-locked neurons at 2 months post-stroke, there was no significant correlation between stroke size and the percentage of time-locked neurons, for either cohort of mice (**Suppl. Fig. S2B**).

Increasing the percentage of C1 whisker-responsive neurons is not the only way that the peri-infarct cortex could contribute to reparative plasticity. For example, we reasoned that, if neurons in barrel cortex that are spared after stroke cannot change their tuning to respond to the whisker that lost its representation, perhaps the original population of C1-responsive neurons in the peri-infarct region could contribute through changes in how they respond to C1 whisker stimulation. For example, forced use therapy could promote recovery by enhancing the magnitude of whisker-evoked activity. To test this, we analyzed sensory-evoked calcium transients for individual neurons in greater detail. Because our stimulation protocol involved 20 sequential whisker stimulations (see Methods), we first aligned calcium signals from all individual epochs of whisker stimuli to generate an average sensory-evoked response for each neuron (**Fig. 6A**). We then calculated both raw z-scores for the magnitude of the calcium transients and the area under the curve (AUC) as measures of magnitude of neuronal responses after whisker stimulation (see Methods). Next, we pooled the average C1 whisker-evoked responses from all time-locked neurons for FOVs within either the C1 barrel or surround barrels (SB) of peri-infarct cortex, for each mouse cohort (whisker-plucked vs. control). For these analyses, we focused on neurons that exhibited time-locked responses to C1 whisker stimulation. C1 whisker stimulation elicited a significant increase in the mean Z-score of calcium fluorescence above baseline in these time-locked neurons (**Fig. 6B**). The average C1 whisker-evoked response at baseline (before stroke) for the control cohort was slightly higher in neurons located within the C1 barrel compared to SB neurons (AUC: 7.65 ± 0.97 vs. 5.86 ± 0.92, respectively) (**Fig. 6B-C**). Importantly, there was no significant difference in Z-scores between plucked and control mice at baseline (**Fig. 6C**).

**Figure 6.**
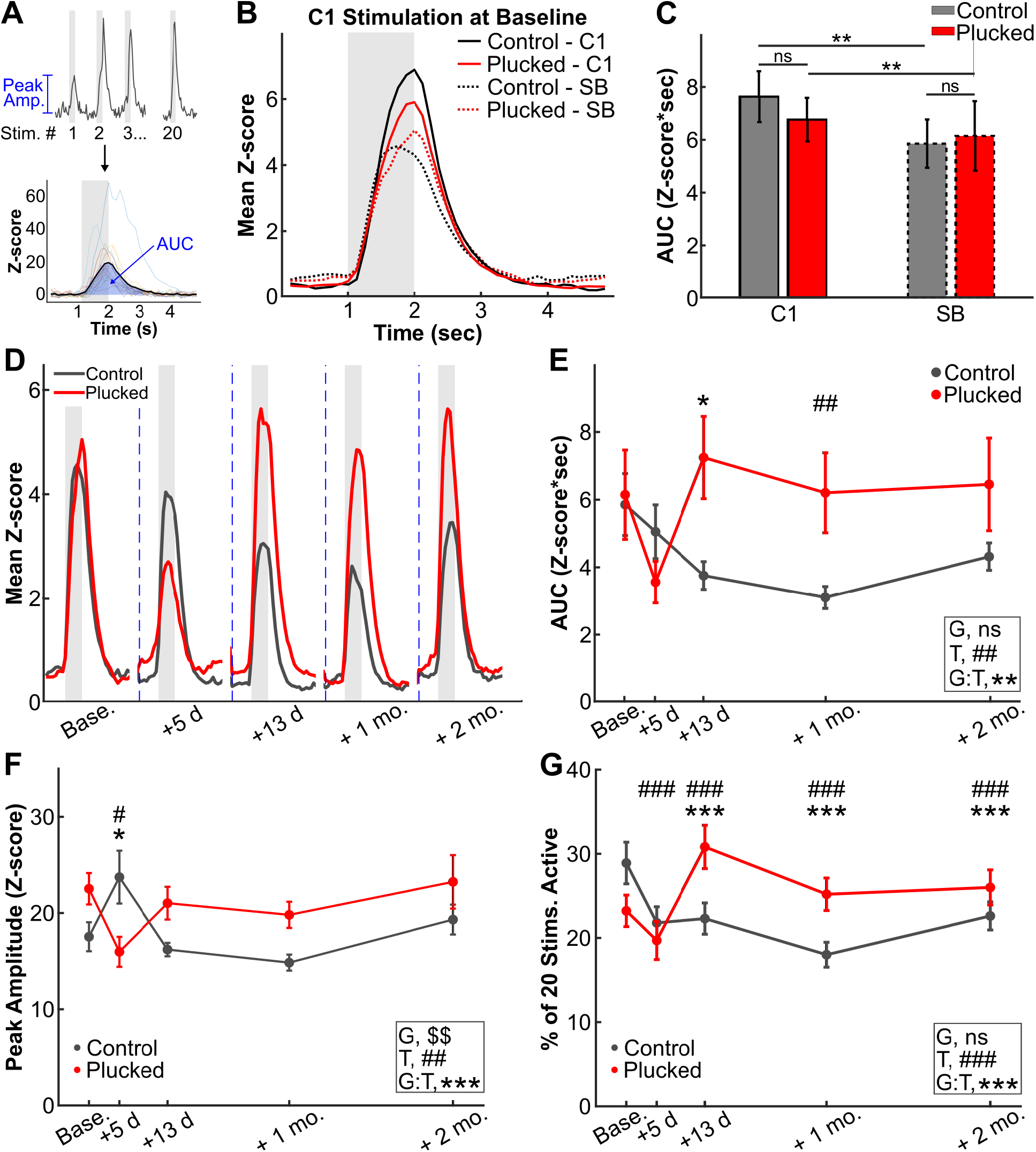
Forced use therapy after stroke increases the reliability of sensory-evoked responses of neurons in peri-infarct cortex. **A**. Analysis pipeline. Neuronal responses to 20 sequential whisker-deflections at 10 Hz (top) were aligned to stimulus onset to generate a mean stimulus-evoked trace per cell (bottom; individual trials are represented by thin lines of different colors; the mean trace is the thick black line). Grey bar denotes stimulus epoch. Peak amplitudes for every stimulus were calculated from individual calcium transients (top), while the area-under-the-curve (AUC) was calculated from the mean stimulus-evoked trace (bottom), see notations in blue. **B.** Mean stimulus-evoked response at baseline (prior to stroke) for all time-locked neurons in control (whiskers intact) vs. forced use mice (all whiskers plucked except C1), comparing neurons located in the C1 barrel (C1; n= 121 neurons from 8 control mice and 145 neurons from 10 plucked mice) to those in surround barrels (SB; n= 128 and 146 neurons from the same mice, respectively). **C.** Quantification of the AUC from the mean stimulus-evoked responses at baseline in *B.* Kruskal-Wallis test, **, *p*<0.01. **D.** Mean stimulus-evoked response from all cells in peri-infarct regions from control or forced use mice over time following stroke. **E.** Quantification of the AUC from the mean stimulus-evoked responses in D. LME model, main effects of group (G, *p*=0.646), timepoint (T, ##, *p*<0.01) and group-by-timepoint interaction (G:T, **, *p*<0.01). Significance for individual coefficients for timepoint (T) or group-by-timepoint interaction (G:T) indicated over corresponding data points (*, *p*<.05; ##, *p*<.01). **F.** Quantification of peak amplitude of stimulus-evoked responses in time-locked neurons over time following stroke, in control vs. forced use (plucked) groups. LME model, main effects of group (G, $$, *p*<0.01), timepoint (T, ##, *p*<0.01) and group-by-timepoint interaction (G:T, ***, *p*<0.001). Significance for individual coefficients for timepoint (T) or group-by-timepoint interaction (G:T) indicated over corresponding data points (* or #, *p*<0.05). **G.** Quantification of the percentage of whisker stimuli that time-locked neurons respond to over time following stroke, in control vs. forced use (plucked) groups. GLME binomial model, main effects of group (G, *p*=0.054), timepoint (T, ###, *p*<0.01) and group-by-timepoint interaction (G:T, ***, *p*<0.001).Significance for individual coefficients for timepoint (T) or group-by-timepoint interaction (G:T) indicated over corresponding data points (*** or ###, *p*<.001).

We next compared the average sensory-evoked response of time-locked cells over time following stroke. In the control stroke group (no rehab), the mean z-score of sensory-evoked responses was significantly lower at 1 month post-stroke, but then gradually increased over 2 months post-stroke, although not quite back to baseline values (**Fig. 6D-E**, black line). In contrast, in the forced use cohort (whisker-plucking), average whisker-evoked responses, which also declined sharply after stroke (3.55 ± 0.62 at +5 d vs. 6.15 ± 1.33 at baseline), subsequently recovered much more rapidly than in the control group starting at 13 d post-stroke (p< 0.01 between groups; **Fig. 6D-E**, red line). Notably, Z-scores remained slightly higher than baseline in the forced use cohort until 2 months post-stroke, though the difference did not reach significance. Similar results were observed when quantifying data from only the first 3 of the 20 stimulation epochs, i.e., those in which neurons are most likely to respond to stimulation, before adaptation to repetitive stimulation is apparent (**Suppl. Fig. 4**).

Average sensory-evoked responses of individual neurons to whisker stimulation could result from either larger amplitude of responses to individual whisker-stimuli, responses to a greater proportion of the whisker stimuli, or both. Therefore, we next quantified responses from time-locked neurons for each of the 20 distinct whisker stimuli delivered during a given imaging session. Peak Z-scores of calcium events differed between groups; after a transient decrease in Z-scores in in the whisker plucked group and an increase in control mice (**Fig. 6F**), by 13 d post-stroke, peak amplitudes had returned to baseline levels in both groups and remained stable through 2 months post-stroke, though they remained slightly higher in the rehab group (**Fig. 6F**). Interestingly, the percentage of stimuli to which individual neurons responded (i.e., had a detectable calcium peak; see Methods), was significantly different between plucked and control cohorts after stroke (**Fig. 6G**); whereas in the control group the percentage of stimuli that neurons responded to remained below baseline levels until 2 months post-stroke, in the whisker-plucked group, it was significantly higher than at baseline (**Fig. 6G**). The difference between groups was greatest at 13 d after stroke, where, in the whisker-plucked group, there was a ~33% increase in the fraction of stimuli that neurons responded to compared to baseline (30.8 ± 2.6 % of stimulation epochs vs. 23.2 ± 1.9 % at baseline), whereas the control group showed a reduction in this value compared to baseline.

We also examined the responses of neurons that were not time-locked to whisker stimulation because we recently found that they are also modulated by whisker stimulation and yet behave differently from time-locked neurons (He et al., 2017). In non-time-locked neurons, there was no detectable increase in mean Z-scores of activity following C1 whisker stimulation, as expected (**Suppl. Fig. 5A**, compare to Fig. 6D). Responses to C1 whisker stimulation in nontime-locked neurons in peri-infarct cortex also changed very little over time following stroke. We found only a slight and transient difference between plucked and control cohorts in C1 whisker-evoked responses for these neurons, as measured by the Z-score of the peak response, at 5 d after stroke (**Suppl. Fig. 5B-C**). The percentage of stimuli for which neurons were active was relatively stable following stroke, with only a small increase seen in neurons from animals in the forced use group (Suppl. **Fig. 5D**). Likewise, when comparing the latency of neurons to reach peak response after whisker stimulation, we found little change over time after stroke in either time-locked neurons or non-time-locked neurons, other than a slight transient decrease at 5 d post-stroke in the control group (**Suppl. Fig. 6**). We also examined the process of adaptation, whereby L2/3 neurons in barrel cortex exhibit progressively smaller responses to ongoing bouts of repetitive whisker stimulation (He et al., 2017). We found that the adaptation index (see Methods) for neurons with time-locked responses to C1 whisker stimulation was similar between the two cohorts of mice (plucked and controls) and did not change significantly over time (**Suppl. Fig. 7**).

Finally, we examined the spontaneous activity (i.e., in the absence of sensory stimulation) of all peri-infarct neurons before and after stroke. Calculating the total AUC of the entire fluorescence trace for each neuron, we found decreased activity in both control and forced use groups at 5 days post-stroke (**Suppl. Fig 8A**). This change recovered to baseline by 13 days post-stroke in the forced use whisker-plucked group, but remained depressed through 2 months post-stroke in the control group. Examining the data more closely, we found small decreases in peak amplitude of spontaneous events in both groups (**Suppl. Fig 8B**) after stroke. More significantly, we found marked changes in frequency of spontaneous calcium transients that paralleled the changes in total AUC (**Suppl. Fig 8C**), specifically, a decreased frequency of calcium transients at 5 days post-stroke that remained below baseline in the control group but recovered (and even increased above baseline) in the forced use group. This result is similar to the effect seen on sensory-evoked responses of time-locked cells after stroke, with the most prominent activity changes found in the likelihood of calcium transients rather than changes in peak amplitude (**Fig. 6E-G**). Taken together, our results show that there is no significant recruitment of new whisker-responsive neurons in the peri-infarct cortex after stroke, either with spontaneous recovery, or with forced use by whisker plucking (**Fig. 7**). Forced use does, however, potentiate the responses of spared whisker-responsive neurons, mainly through increased likelihood of response to sensory stimuli.

**Figure 7.**
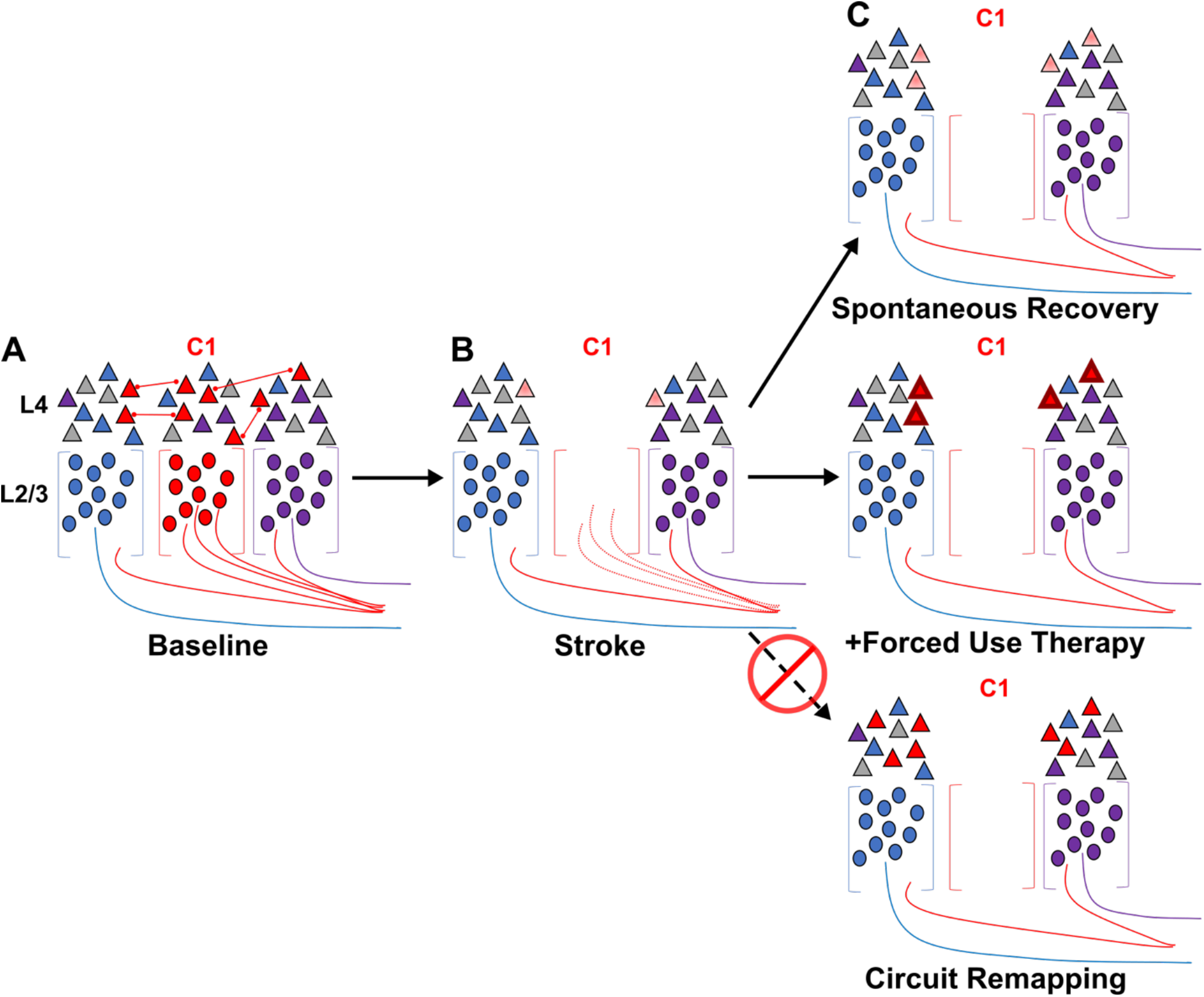
Models of L2/3 whisker somatosensory circuit changes post stroke. **A**. Distribution of whisker responsive L2/3 pyramidal neurons (triangles) in 3 adjacent barrels of the S1BF at baseline. Colors denote responsivity to a particular whisker (red = C1). Thalamocortical axons are depicted innervating L4 stellate neurons (circles) within barrels, with barrel septa pictured as brackets. Intracortical connections between similarly tuned L2/3 pyramidal neurons are depicted as well. Note that within a given barrel, L2/3 neurons are tuned to both the principal and surround whiskers. **B.** After stroke targeting a single barrel, all neurons within the barrel are destroyed. The proportion of surround barrel neurons that are tuned to the C1 whisker (corresponding to the infarcted barrel) is decreased after stroke and their sensory-evoked responses are reduced (paler shading). **C.** Spontaneous recovery (top panel) results in restoration of the proportion of surround barrel neurons tuned to the C1 whisker destroyed by stroke, but their sensory-evoked responses remain depressed, and there is no re-tuning of spared neurons to replace neurons lost to stroke. Forced use therapy (middle panel) after stroke restores and potentiates sensory-evoked responses in surround whisker neurons, but does not result in true circuit remapping with recruitment of new C1 whisker-responsive neurons (bottom panel).

## Discussion

A major gap in our understanding of functional recovery after stroke has been the lack of clarity regarding the specific changes in neuronal activity that mediate such recovery. The prevailing dogma in stroke is that circuits can remap by recruiting surviving neurons to assume new functions previously encoded by neurons that died (Murphy and Corbett, 2009; Winship and Murphy, 2009). Although this hypothesis is attractive and makes intuitive sense, the evidence to support it has been largely indirect and ultimately unconvincing (see below). Here, we sought to test the remapping model by recording neural activity in peri-infarct cortex before and after a small single barrel stroke. Using three different in vivo approaches to identify remapped circuits (ISI, 2P calcium imaging and TRAP), we find no evidence of remapping of lost functionalities. That is, we could not identify any increase in the population of C1 whisker-responsive neurons that would be expected if surviving neurons were multitasking to assume the role of the dead neurons. On the contrary, in animals allowed to recover spontaneously, we find that the proportion of whisker-responsive cells decreases acutely after stroke before returning only to baseline levels over 1-2 months. However, in rehabilitated animals (using whisker plucking as a means of forced use therapy), we find significant increases in the reliability of whisker-evoked responses in circuits that were already responding to that whisker before the stroke. Our findings are significant because they put into question the long-held remapping model of stroke recovery that has influenced stroke research for decades.

### Overcoming limitations of previous studies that support the remapping theory

Much of the remapping hypothesis for stroke recovery is predicated on human brain mapping studies documenting examples of reorganization of cortical sensorimotor activity maps post-stroke (Altamura et al., 2007; Carey et al., 2011; Cramer et al., 1997; Delvaux et al., 2003; Jang, 2011; Jang et al., 2005; Rossini et al., 1998; Schaechter et al., 2006; Zemke et al., 2003). Because these studies revealed a number of different mechanisms for circuit rearrangements after stroke, they failed to yield a unifying theory of remapping. Indeed, there is significant inherent variability in those studies, including lesion location, lesion size, or time of imaging after stroke (acute vs. chronic). Those imaging methods also suffer from several technical and practical limitations. For example, functional fMRI and fluorodeoxyglucose PET rely on surrogate markers of neuronal activity, such as blood flow, oxygen content or metabolism (all of which are altered by stroke). These approaches are also missing a pre-stroke baseline imaging session, which would be critical to interpret whether remapping actually occurred or whether observed changes just represent baseline variability across subjects. Most importantly, these techniques lack the temporal and spatial resolution necessary for recording activity in single neurons over time (Crosson et al., 2010). Our goal was precisely to overcome these limitations, by recording from individual neurons or populations longitudinally before and after stroke, using a single barrel stroke model that leads to consistent and reproducible infarcts.

### No evidence of remapping after single barrel strokes

In order to test the remapping hypothesis, we reasoned the S1BF would be an ideal location because it exhibits strong somatotopic organization, with each individual whisker being primarily mapped to a single cortical column, or barrel (Diamond and Arabzadeh, 2013). However, within an individual barrel, neurons exhibit heterogenous tuning, with a plurality of neurons tuned to the corresponding principal whisker, but many others tuned to surround whiskers (Clancy et al., 2015). Such tuning is highly dynamic in the intact brain, with neurons capable of modifying their selectivity in response to changes in sensory experience. For example, after whisker plucking, a well-established paradigm of peripheral sensory deprivation, individual L2/3 neurons rapidly change their tuning away from deprived whiskers and toward spared whiskers (Margolis et al., 2012). Such circuit plasticity can be observed at a macroscopic level as well, with expansion of spared whisker cortical activity maps using 2-deoxyglucose or ISI mapping (Gao et al., 2017; Polley et al., 1999). However, whether lesioning the central whisker representation in the cortex, i.e., its corresponding barrel, triggers similar changes in tuning of neurons in surrounding barrels has never been documented.

After targeting PT strokes to a single barrel (C1) in S1BF we failed to see any evidence of remapping to neighboring regions using ISI in the majority of mice, even up to 1-2 months after stroke. A minority of mice had smaller, weaker maps in the approximate location of the original C1 map, suggesting perhaps incomplete ablation of the entire C1 barrel. Our results are consistent with a prior study that found smaller and weaker ISI maps following forelimb somatosensory PT strokes (Tennant et al., 2017). Using calcium imaging in the D3 barrel adjacent to the C1 barrel, we found that a minority of L2/3 neurons responded to C1 whisker stimulation at baseline, as reported previously (Clancy et al., 2015). Given the presence of these surround whisker-tuned neurons, we expected to find that after a stroke targeting the C1 barrel we would identify spared neurons in surround barrels that changed their tuning, resulting in greater percentages of C1 whisker-responsive neurons; but instead we found the opposite result. Confirming this, we also did not find an increase in putative C1 whisker-responsive neurons using activity-dependent labeling in TRAP mice, nor in our separate forced use therapy cohort. Overall, our data provide robust evidence against the remapping hypothesis for stroke recovery: there is no new recruitment of spared neurons to subsume lost functionalities.

### What about the prior animal studies supporting the remapping hypothesis?

A second argument for the remapping hypothesis for stroke recovery has come from molecular and structural imaging in animal models that demonstrated changes consistent with neural plasticity. These include transcriptomic changes involving growth factors and plasticity related genes (Carmichael et al., 2005), sprouting of axons (Li et al., 2010), and changes in dendritic spine turnover (Brown et al., 2007; Clark et al., 2019; Mostany et al., 2010). However, definitive evidence showing that these changes actually result in neurons or circuits assuming the functions of neurons that were lost to stroke remains lacking. Transcriptional changes could support structural plasticity, such as dendritic spine turnover or axonal branching, but this may reflect circuit plasticity to augment existing connectivity rather than functional remapping. Indeed, structural plasticity triggered by loss of presynaptic boutons (from loss of neurons in the infarct) would likely lead to new connections that enhance pre-existing connections rather than create new ones.

Perhaps the best evidence to date for the remapping theory as a general mechanism for functional recovery after stroke was the pioneering study of Winship and Murphy in 2008, in which they reported that a small but significant population of neurons exhibited broader tuning 1-2 months post-stroke; for example, neurons in hindlimb S1 also responded to contralateral forelimb (FL) stimulation, or neurons in FLS1 responded to both contralateral and ipsilateral FL stimulation (Winship and Murphy, 2008). There are several important differences between that study and ours that may account for our discrepant findings, including the stroke location and brain region imaged (FLS1or HLS1 vs. S1BF), and the type of imaging (acute recordings with Oregon Green BAPTA-1 in different mice before and after stroke vs. chronic imaging of GCaMP6s neurons in the same mice over time). Although we are not aware of other studies recording sensory-evoked activity of cortical neurons in the peri-infarct cortex, the activity of FL thalamocortical axons is persistently impaired post-stroke (Tennant et al., 2017), a result which is generally in line with our findings. Even if remapping were to occur, it remains unclear what the functional consequences of broadened tuning of neurons post-stroke might be. Sparse firing of cortical neurons to specific stimuli is thought to underlie efficient sensory processing in S1 (Petersen and Crochet, 2013). One could imagine that, as spared neurons assume additional roles post-stroke, this might degrade sensory processing. In line with this hypothesis, we and others found that even a small increase in tuning width of neurons in the visual cortex leads to impaired visual discrimination in mice (Goel et al., 2018; Lee et al., 2012).

### Potential limitations of our study

One could argue that there are reasons why we failed to detect any remapping. One possibility is that we were imaging in the wrong location, as we surveyed regions surrounding the C1 barrel rather than more distant locations in S1 or even subcortical regions. However, this is unlikely because the tuning of L2/3 neurons in adjacent of barrels is known to be promiscuous (Clancy et al., 2015) and plasticity triggered by whisker plucking in the intact S1BF preferentially recruits adjacent barrels (Glazewski and Fox, 1996; Margolis et al., 2012). Subcortical plasticity may occur, especially at the level of the thalamus, but this is probably insufficient to completely restore sensory function; it may help with whisker detection, but for discrimination, S1BF is essential (Miyashita and Feldman, 2013; Park et al., 2020). Another possibility is that our strokes were too small to produce a sufficient functional deficit to drive remapping. However, other studies have shown that single barrel ablation does lead to functional impairment (Shih et al., 2013). We also found that larger strokes were actually associated with a trend toward even fewer whisker-responsive cells at 2 months. Finally, it is possible that our calcium imaging approach was not sensitive enough to detect remapping. Yet, we were able to detect significant increases in the reliability of how surviving neurons respond to their principal whisker, and that whisker plucking further enhanced such changes, arguing against this possibility. Thus, plasticity does occur after stroke, but it just does not appear to involve remapping. It is also unlikely that remapping is mediated by a small number of neurons scattered throughout peri-infarct cortex because we actually probed dozens of whisker-responsive neurons in vivo and found only one animal (out of 18) with a significantly higher percentage of neurons that responded to the C1 whisker after stroke, compared to baseline.

### Maladaptive plasticity in peri-infarct cortex

The notion that stroke triggers changes that actually hamper, rather than promote, recovery (maladaptive plasticity) has been suggested previously. The fact that whisker plucking (which triggers tuning changes in healthy S1 cortex) did not promote circuit remapping after stroke suggests that mechanisms exist within the peri-infarct cortex that limit the potential for remapping. This interpretation is supported by prior studies showing impaired plasticity in peri-infarct cortex, including the absence of whisker trimming-induced expansion of cortical maps (Jablonka et al., 2007, 2012), and loss of monocular deprivation-induced ocular dominance shifts (Greifzu et al., 2011). The mechanisms underlying this impairment in plasticity are not known, but may include alterations in the excitation/inhibition balance, such as increased tonic inhibition (Alia et al., 2016; Clarkson et al., 2010; Hiu et al., 2016). The fact that C1 whisker-evoked responses were reduced after stroke could be consistent with such an increase in inhibition. Given that whisker plucking is known to induce circuit remapping via disinhibition in the intact S1BF, we favor a model in which increased inhibition limits circuit remapping after stroke, though further studies will be necessary to clarify these mechanisms.

### Forced used therapy potentiates the responses of surviving neurons but does not induce remapping

Although we did not find that forced use therapy induced circuit remapping after stroke, we did observe that it potentiated whisker-evoked responses in spared C1 responsive neurons, driven largely by a significant increase in the reliability of responses (**Fig. 6G**). We may have identified the circuit basis for how rehabilitation via constraint-induced movement therapy helps the recovery of stroke patients. Specifically, we hypothesize that this rehabilitation technique works by boosting responses of spared SB neurons that were already responding to the C1 whisker before stroke, rather than by changing the selectivity of other neurons so that they become newly tuned to the C1 whisker. More broadly, our results have significant implications for physical rehabilitation strategies, as these are often designed to engage neuroplasticity mechanisms known to be operant in healthy brain (Di Pino et al., 2014), but which our results, and others, suggest may be impaired post-stroke. In order to promote true circuit remapping (**Fig. 7C**, right panel), rehabilitation may need to be combined with genetic (Caracciolo et al., 2018), pharmacologic (Clarkson et al., 2010), or other approaches (Cheng et al., 2016; Tennant et al., 2017) to circumvent mechanisms limiting remapping post stroke.

In conclusion, while the lack of remapping after stroke we describe is unexpected and perhaps counterintuitive to some, it cannot be ignored. Functional recovery, if and when it occurs after stroke, is not mediated by surviving neurons assuming new roles (multi-tasking) but may instead involve allocating resources to potentiate pre-existing circuits, at least in somatosensory cortex. Until it is unequivocally demonstrated that neurons can consistently undertake the roles of neighbors that were lost after cortical lesions, the notion of circuit remapping after stroke should be viewed with skepticism.

## Acknowledgments

This work was supported by R01NS076942-05 and R25-NS065723 from NIH-NINDS. We thank Drs. Cynthia He and Daniel Cantu for MATLAB analysis code (which was adapted for this work), Drs. Chi Hong Tseng and Sitaram Vangala for advice on statistical analysis using mixed effects models, Dr. Ricardo Mostany and Grant Higerd for their early ISI experiments in stroke, and Drs. Anubhuti Goel, Nazim Kourdougli, Anand Suresh, and Bo Wang for helpful discussions.

## Supplementary Figures

**Supplementary Figure 1.**
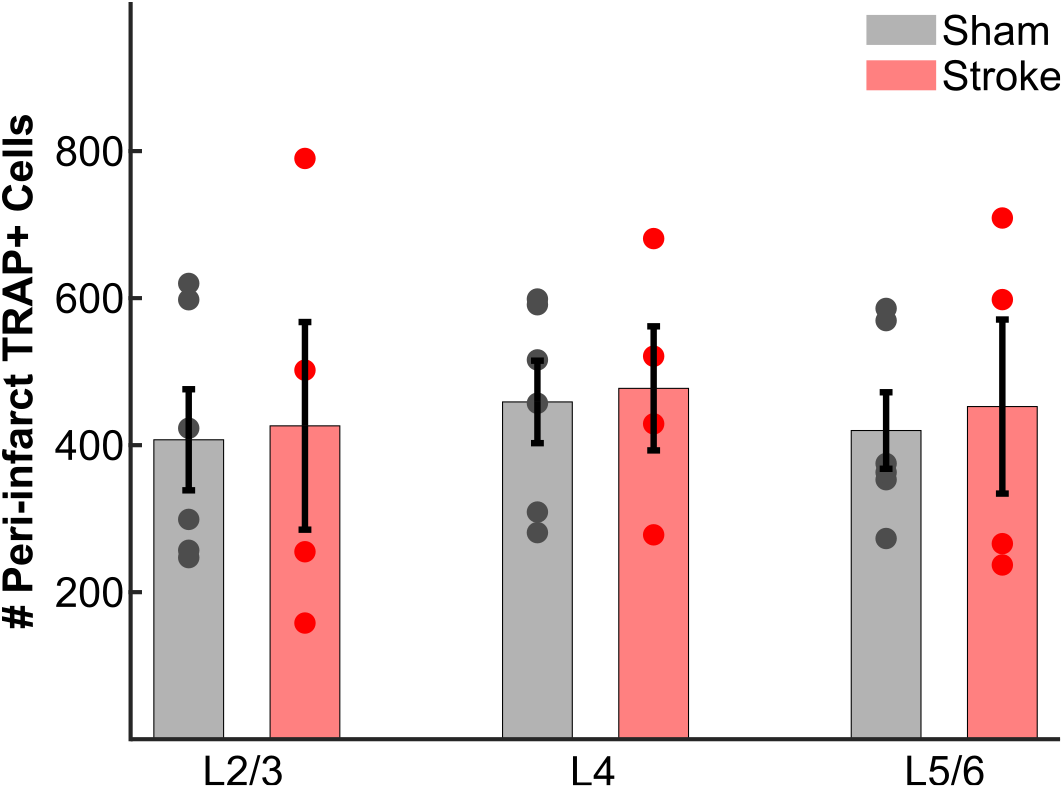
TRAP labeling of C1 whisker-responsive neurons in the peri-infarct cortex by cortical layers. Quantification of the total number of TRAP+ neurons in the peri-infarct cortex (stroke group) or surround barrels (sham group), separated by cortical layers. No significant differences were found using a two-way ANOVA for effects of Group (*p*=0.736), Layer (*p*=0.828), or Group*Layer (*p*=0.995).

**Supplementary Figure 2.**
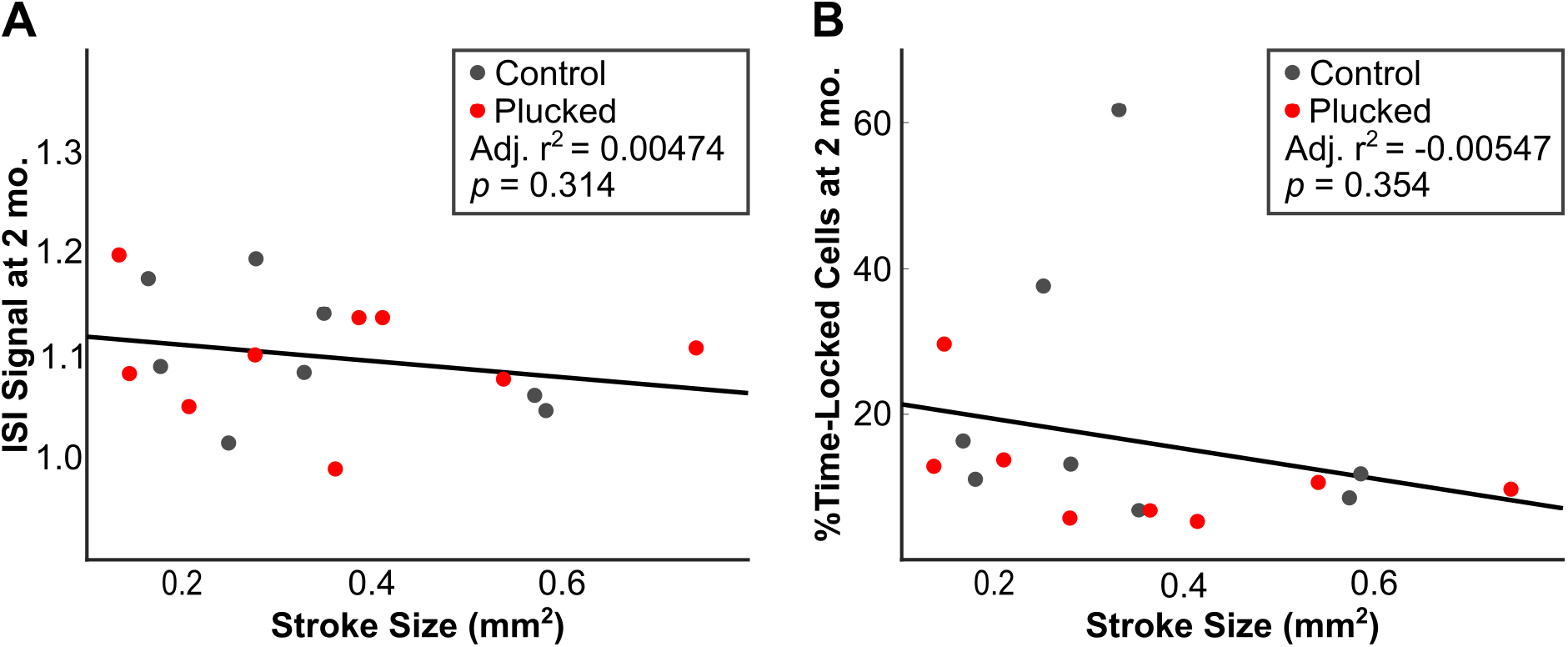
No significant correlation between stroke size and C1 whisker evoked activity. **A.** Correlation between C1-whisker evoked ISI signal at 2 months with stroke size at Day 5. The plotted line represents the correlation for pooled data from both control and forced use (plucked) groups. **B.** Correlation between percentage of neurons with time-locked responses in peri-infarct regions at 2 months with stroke size at Day 5. The plotted line represents the correlation for pooled data from both control and forced-use (plucked) groups.

**Supplementary Figure 3.**
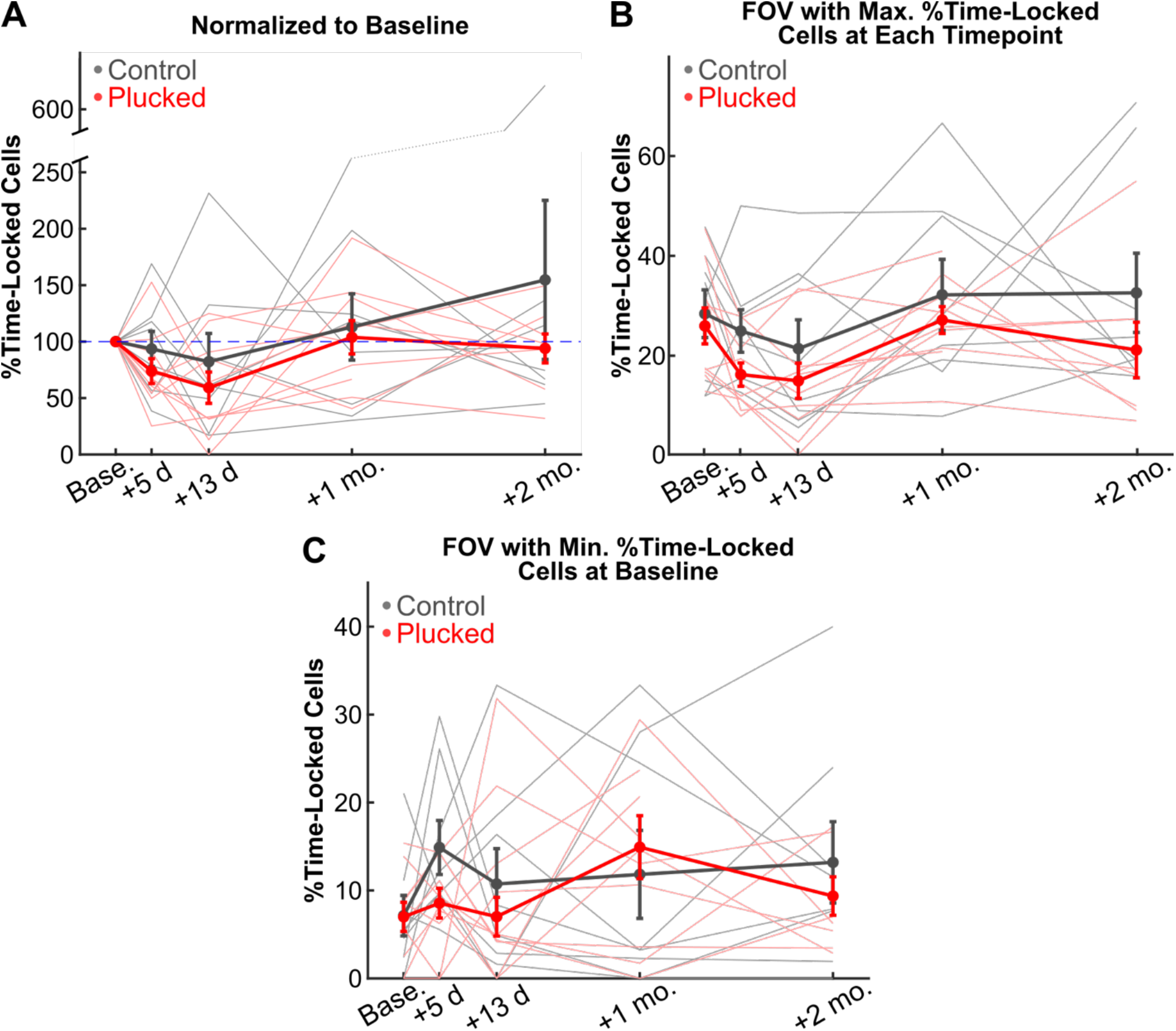
Trends for the change in the number of C1-whisker time-locked cells after stroke with forced-use therapy are similar across multiple methods of analysis. **A.** Percentage of neurons in peri-infarct regions across all FOV with responses that are time-locked to C1-whisker stimulation normalized to the baseline percentage of time-locked cells in the peri-infarct regions. **B.** Percentage of neurons in peri-infarct cortex with responses that are time-locked to C1-whisker evoked activity when analysis is restricted to the FOV with the greatest number of time-locked cells for each mouse at each timepoint. **C.** Percentage of neurons in peri-infarct cortex with responses that are time-locked to C1-whisker evoked activity when analysis is restricted to the FOV with the least number of time-locked cells at baseline for each mouse (to test whether peri-infarct regions with the fewest time-locked cells to the C1 whisker might be more capable of assuming that role post-stroke; the answer is clearly no).

**Supplementary Figure 4.**
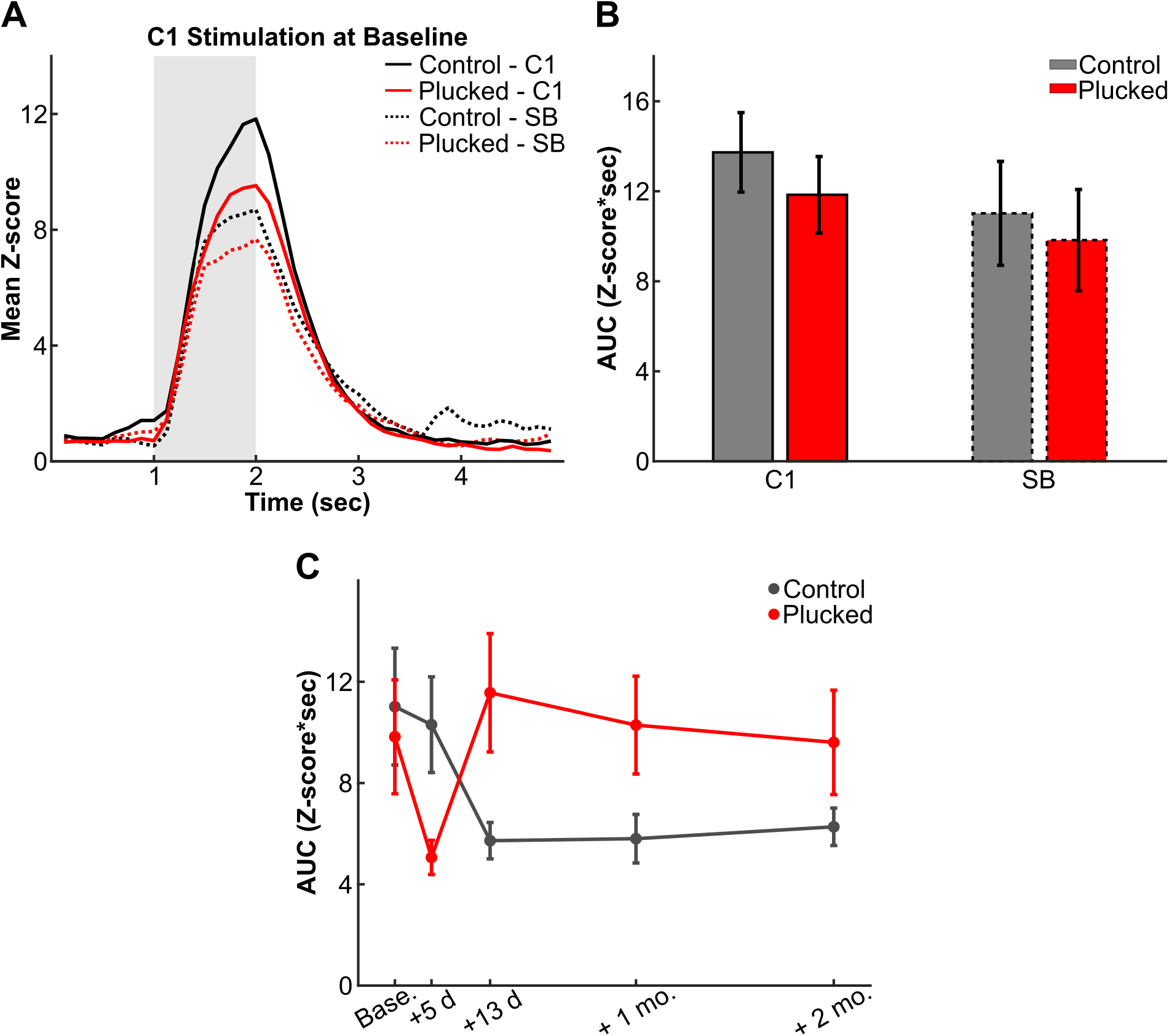
Sensory evoked responses of time-locked neurons (analysis restricted to the first 3 of 20 stimulation epochs, to control for adaptation). **A.** Mean stimulus-evoked response (calculated from the first 3 of 20 stimulation epochs) at baseline (prior to stroke) for all time-locked neurons in control (whiskers intact) or forced use (all whiskers plucked except C1) mice, comparing neurons located in the C1 barrel (C1) to those in surround barrels (SB). **B.** Quantification of the AUC from the mean stimulus-evoked responses at baseline in A. **C.** Quantification of the AUC from the mean stimulus-evoked responses from time-locked neurons in peri-infarct surround barrels over time after stroke.

**Supplementary Figure 5.**
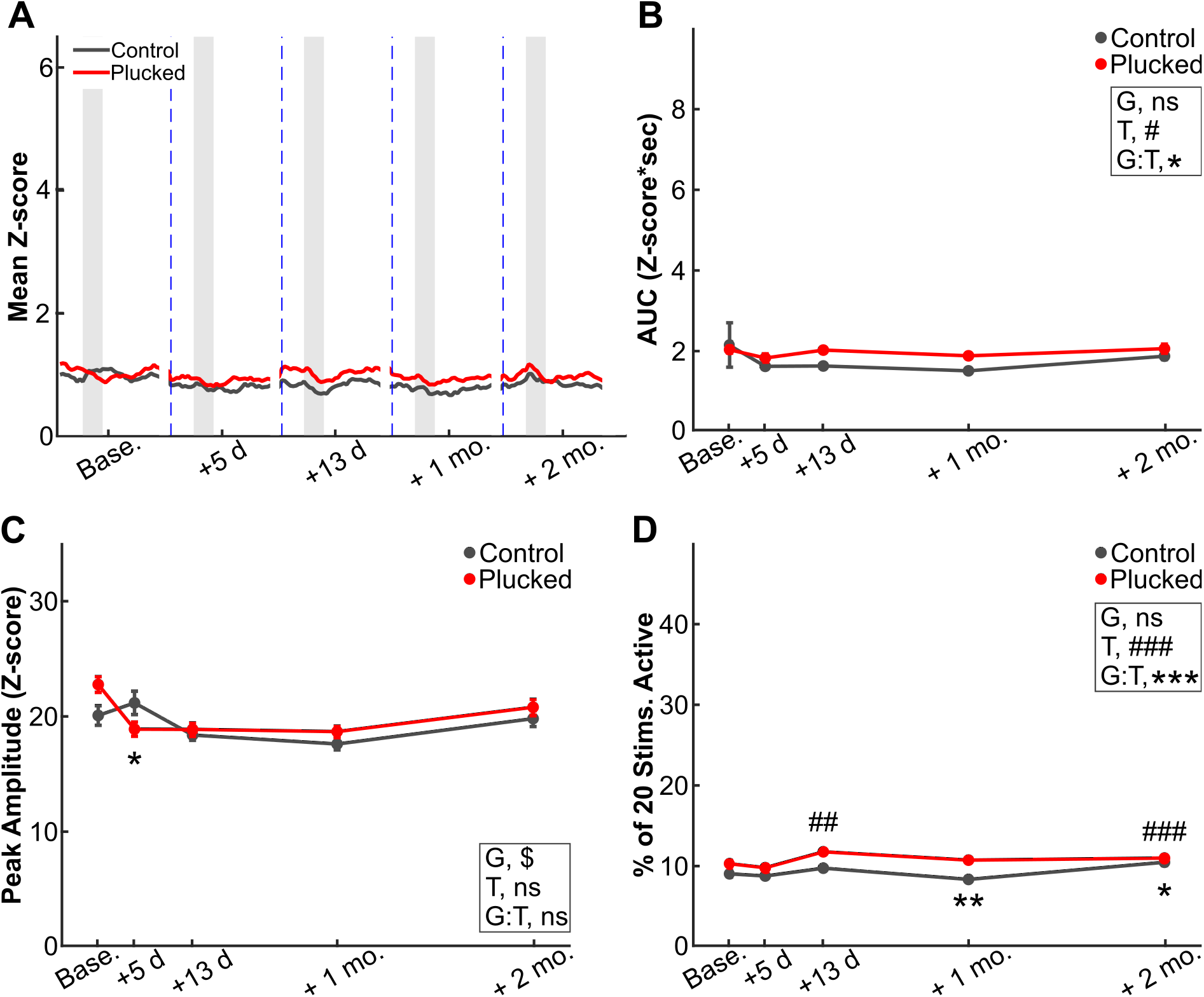
Sensory-evoked responses in neurons in peri-infarct cortex that were not time-locked to whisker stimulation (data plotted with same axes as in Fig. 6). **A.** Mean stimulus-evoked response from non time-locked cells in control or forced use (plucked) groups in peri-infarct regions over time following stroke. **B.** Quantification of the AUC from the mean stimulus-evoked responses in A. LME model, main effects of group (G, *p*=0.065), timepoint (T, #, *p*<0.05) and group-by-timepoint interaction (G:T, *, *p*<0.05). None of the individual coefficients for timepoint (T) or group-by-timepoint interaction (G:T) were significant. **C,D.** Quantification of peak amplitude (C) and fraction of whisker stimuli that neurons respond to (D) for time-locked cells over time following stroke in control and forced use (plucked) groups. Mixed effects models, main effects of group (G), timepoint (T) and group-by-timepoint interaction (G:T) are indicated on the respective plots. Significance for individual coefficients for timepoint (T) or group-by-timepoint interaction (G:T) indicated under corresponding data points (* or #, *p*<0.05; ** or ##, *p*<0.01; *** or ###, *p*<0.001).

**Supplementary Figure 6.**
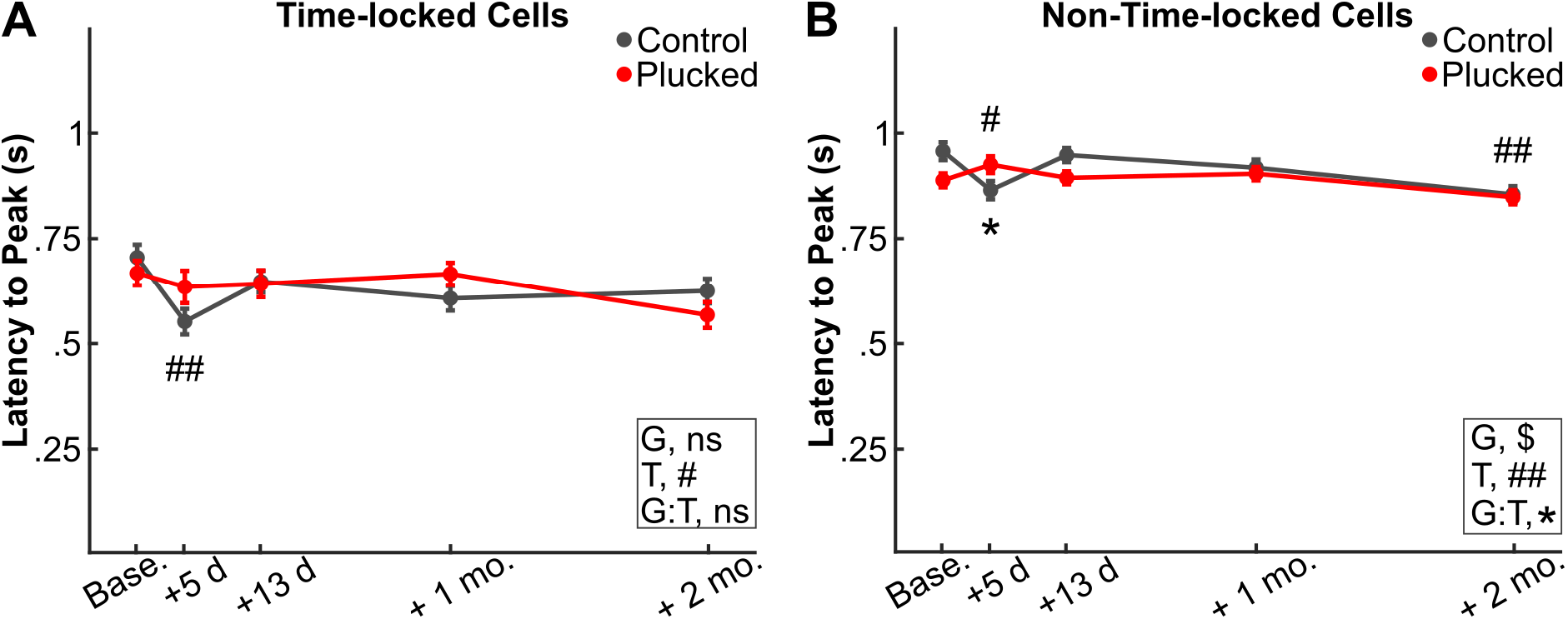
Latency to sensory-evoked peak amplitude is not affected by stroke or forced use. **A,B.** Quantification of latency to peak amplitude from whisker stimulus onset in neurons time-locked (A) or non-time-locked neurons (B) over time following stroke in control and forced use (plucked) groups. Mixed effects models, main effects of group (G), timepoint (T) and group-by-timepoint interaction (G:T) are indicated on the respective plots. Significance for individual coefficients for timepoint (T) or group-by-timepoint interaction (G:T) indicated under corresponding data points (* or #, *p*<0.05; ** or ##, *p*<0.01).

**Supplementary Figure 7.**
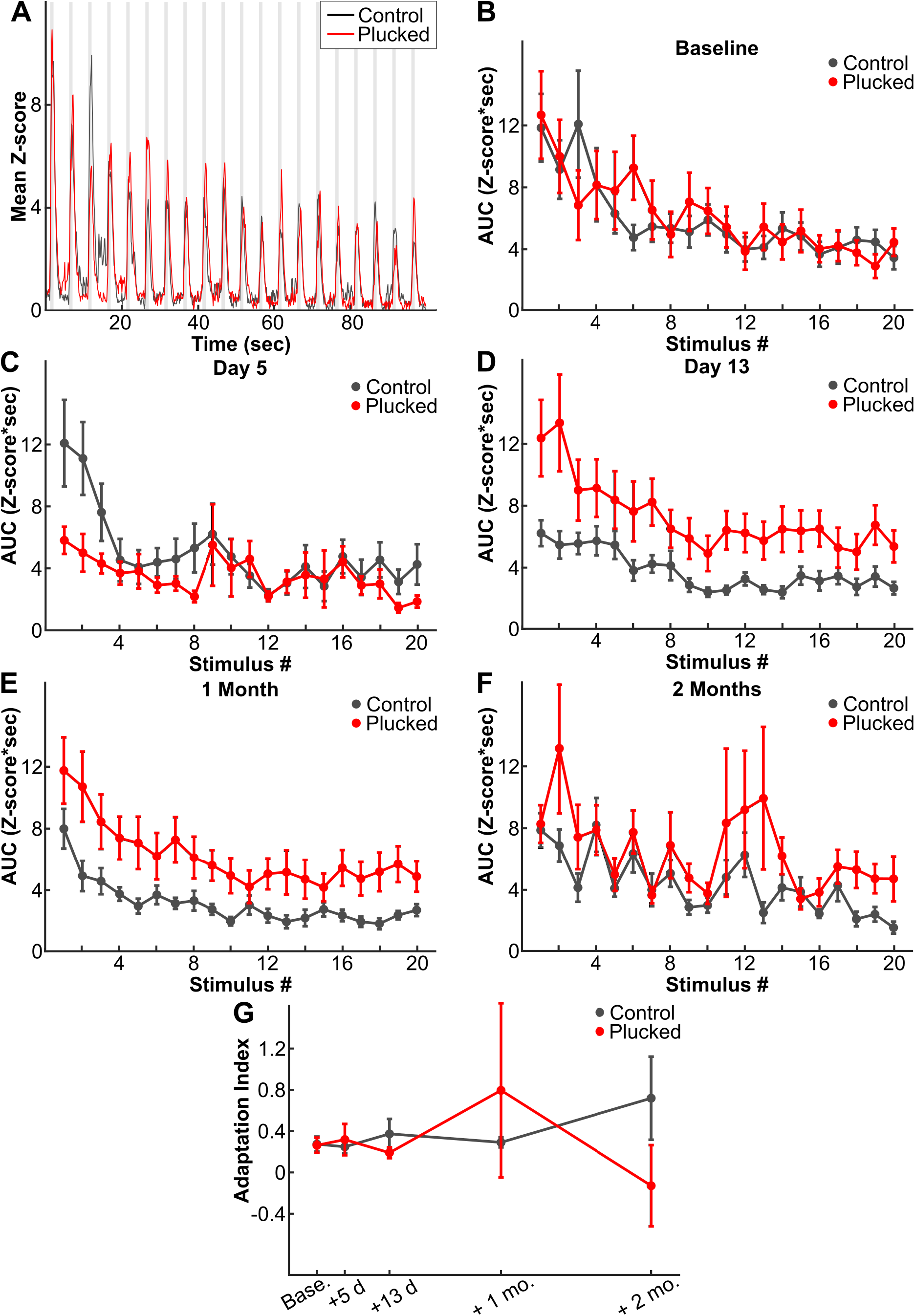
Neuronal adaptation to repetitive whisker stimulation in peri-infarct cortex. **A**. Mean Z-score of cells from control or forced-use groups at baseline, aligned to onset of whisker stimulus, for each of 20 consecutive stimuli. **B-F.** Quantification of mean AUC of sensory-evoked responses from control or forced-use groups at baseline (B), +5 days (C), +13 days (D), +1 month (E), or +2 months (F), aligned to onset of whisker stimulus, for each of 20 consecutive stimuli. **G.** Adaptation index for sensory-evoked responses in peri-infarct cortex over time following stroke. LME model, main effects of group (G, *p*=0.983), timepoint (T, *p*=0.903) and group-by-timepoint interaction (G:T, *p*=0.471).

**Supplementary Figure 8.**
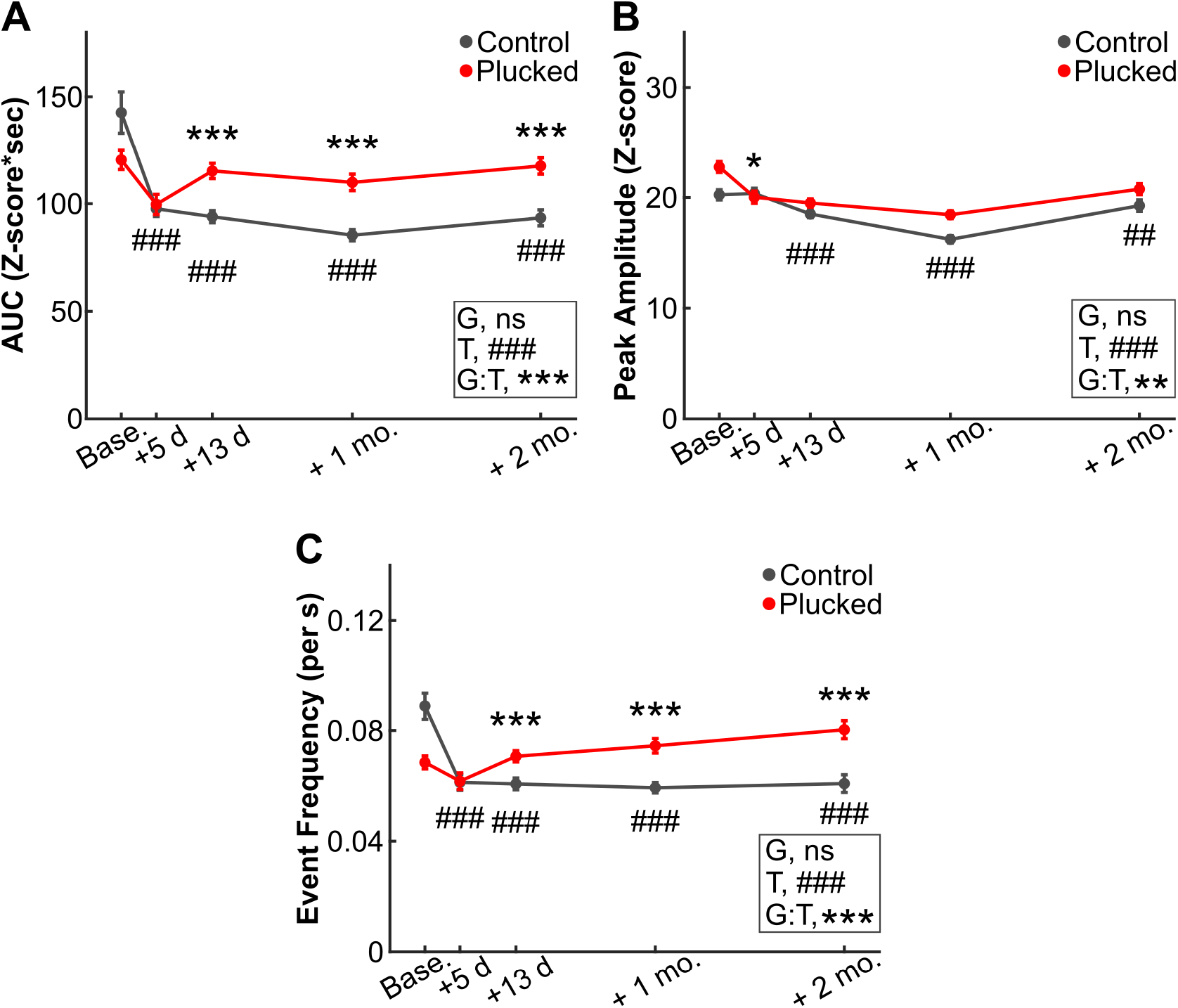
Spontaneous activity of peri-infarct neurons over time before and after stroke. **A.** Quantification of the AUC from the entire fluorescence trace (~100 s) of spontaneous activity from peri-infarct neurons in control and forced-use (plucked) groups before and after stroke. **B, C.** Quantification of peak amplitude (B) and frequency (C) of calcium events for peri-infarct neurons in control and forced-use (plucked) groups before and after stroke. Statistics for all plots from LME models, with main effects of group (G), timepoint (T) and group-by-timepoint interaction (G:T) indicated on the respective plots. Significance for individual coefficients for timepoint (T) or group-by-timepoint interaction (G:T) are indicated under corresponding data points (* or #, *p*<0.05; ** or ##, *p*<0.01; *** or ###, *p*<0.001).

## Notes

### Competing Interest Statement

The authors have declared no competing interest.

